# Characterization of *mWake* expression in the murine brain

**DOI:** 10.1101/2020.05.25.114363

**Authors:** Benjamin J. Bell, Annette A. Wang, Dong Won Kim, Seth Blackshaw, Mark N. Wu

**Affiliations:** McKusick-Nathans Department of Genetic Medicine, Johns Hopkins University, Baltimore, MD 21287; Department of Neurology, Johns Hopkins University, Baltimore, MD 21205; Department of Neuroscience, Johns Hopkins University, Baltimore, MD 21205

**Author notes:** These authors contributed equally (Benjamin J. Bell, Annette A. Wang). Correspondence should be addressed to M.N.W.

**Keywords:** mWAKE, WAKE, circadian, hypothalamus, suprachiasmatic nucleus

## Abstract

Structure-function analyses of the mammalian brain have historically relied on anatomically-based approaches. In these investigations, physical, chemical, or electrolytic lesions of anatomical structures are applied, and the resulting behavioral or physiological responses assayed. An alternative approach is to focus on the expression pattern of a molecule whose function has been characterized and then use genetic intersectional methods to optogenetically or chemogenetically manipulate distinct circuits. We previously identified WIDE AWAKE (WAKE) in *Drosophila*, a clock output molecule that mediates the temporal regulation of sleep onset and sleep maintenance. More recently, we have studied the mouse homolog, mWAKE/ANKFN1, and found that its role in the circadian regulation of arousal is conserved. Here, we perform a systematic analysis of the expression pattern of *mWake* mRNA, protein, and cells throughout the adult mouse brain. We find that mWAKE labels neurons in a restricted, but distributed manner, in multiple regions of the hypothalamus (including the suprachiasmatic nucleus), the limbic system, sensory processing nuclei, and additional specific brainstem, subcortical, and cortical areas. Interestingly, mWAKE is also observed in non-neuronal ependymal cells. In addition, to describe the molecular identities and clustering of *mWake*^+^ cells, we provide detailed analyses of single cell RNA sequencing data from the hypothalamus, a region with particularly significant mWAKE expression. These findings lay the groundwork for future studies into the potential role of *mWake^+^* cells in the rhythmic control of diverse behaviors and physiological processes.

## Introduction

For centuries, scientists have endeavored to attribute aspects of cognition, emotion, and behavior to anatomic regions of the brain (Grand, 1999). Initial efforts involved resecting or lesioning a brain area and observing its impact on a specific behavior. For example, ablation studies in the 19^th^ century revealed the importance of the medulla in respiratory control and of the frontal lobes for attention (Bianchi, 1895; Pearce, 2008). Contemporary investigations have refined this classical anatomical approach by utilizing cell type-specific ablations (Nirenberg & Cepko, 1993). However, an alternative method for delineating structure-function relationships starts from characterizing the roles of specific molecules and then examining their expression pattern to identify neural circuits involved in relevant behaviors. For example, in *Drosophila*, *fruitless* and *doublesex*, which encode critical factors in sex determination, mark neural circuits involved in a variety of sexually dimorphic behaviors (Auer & Benton, 2016). Similarly, in mammals, neurons expressing the androgen receptor label sexually dimorphic circuits in the preoptic area (POA) and basal forebrain (BF) (Shah et al., 2004).

From a forward-genetic screen in *Drosophila*, we previously identified WIDE AWAKE (WAKE), a molecule that regulates the circadian timing of sleep (Liu et al., 2014). WAKE acts downstream of the circadian clock to promote sleep during the night, by inhibiting the excitability of arousal-promoting clock neurons at that time (Liu et al., 2014). This protein has a single mammalian ortholog, mWAKE/ANKFN1/NMF9, which is enriched in the suprachiasmatic nucleus (SCN), the circadian pacemaker in mammals (Liu et al., 2014; Zhang et al., 2015). Our recent work on mWAKE suggests that the function of WAKE in regulating arousal by rhythmically modulating neural activity is conserved in mice (Bell et al., 2020).

Here, in an effort to identify additional brain regions whose activity and related behaviors depend on mWAKE and the circadian clock, we use three independent techniques to investigate mWAKE expression throughout the mouse brain. We use the highly sensitive RNAscope *in situ* hybridization (ISH) technique to label *mWake* mRNA and also perform immunofluorescence (IF) and imaging studies from two different transgenic mouse models to visualize mWAKE protein and mWAKE^+^ neuronal processes. To gain a deeper understanding of the molecular identities of *mWake^+^* cells and how they form different sub-clusters within an anatomical region, we also performed single cell RNA sequencing (scRNA-Seq) analyses on *mWake^+^* cells in the hypothalamus. Together, these investigations revealed that mWAKE is expressed in a restricted manner across multiple regions of the mouse brain. These areas span the brainstem, subcortical areas, and cortex, with particularly prominent expression in the hypothalamus, and include not only *mWake*^+^ neurons but also ependymal cells that line the cerebral ventricles. Although the precise functions of these *mWake*^+^ regions remain to be determined, their locations are suggestive of potential roles in circadian-related behaviors, arousal, sensory processing, and emotion. These results comprise a catalog of *mWake* expression in the mouse brain and *mWake*^+^ cell identity in the hypothalamus and may ultimately lead to the identification of neural circuits mediating the circadian regulation of arousal and other internal states and behaviors.

## Experimental design

### Animals

All experiments and animal procedures were approved by the Johns Hopkins Institutional Animal Care and Use Committee (IACUC). Animals were raised in a common animal facility, group housed and maintained with food and water *ad libitum*. Male mice were used in all experiments, at 2-4 months old for all histology and 6-7 weeks of age for scRNA-Seq. All genotyping was performed via Taq-Man based rtPCR probes (Transnetyx). Wild-type (WT) mice in all experiments were *C57BL/6J. WAKE^Cre^* mice carry an insertion of a tdTomato-p2A-Cre cassette in exon 5 and was previously described in Bell et al., 2020 (Bell et al., 2020).

#### mWAKE^V5^ mice

A V5 (GKPIPNPLLGLDST) epitope tag was fused to the C-terminus of mWAKE directly before the stop codon using CRISPR/Cas-9 genome editing (Hsu, Lander, & Zhang, 2014), with a gRNA (TTA AGT AGT ATG CTT TAG GG) targeting the stop codon of *mWake* and a 150 bp oligonucleotide (AGT TCA GAG ATG AGT CCA GAC CCC ACA TCT CCA GTT TCA GAA ATA TTA AGT AGT ATG CTT GGC AAG CCC ATC CCC AAC CCC CTG CTG GGC CTG GAC AGC ACC TAG GGT GGC CCA CAC CGG CTC TCT ATT TGT CCC CTG CTA TTC CTT GCA TTT CTT CAG CAC AGC) for homology-directed repair to insert the V5 polypeptide sequence. mWAKE-V5 was then detected using immunofluorescence staining in fixed tissue sections. *mWAKE^V5^* was backcrossed to the *C57BL/6J* background five generations before being used in any experiments.

### Immunohistochemistry

Mice were deeply anesthetized with a ketamine/xylazine mixture and then fixed by transcardial perfusion with 4% paraformaldehyde. Brains were subsequently drop-fixed in 4% paraformaldehyde for 24-48 hrs and transferred into 1xPBS (137 mM NaCl, 2.7 nM KCl, 10 mM Na_2_HPO_4_, 1.8 nM KH_2_PO_4_) before being sectioned at 40 μm thickness using a VT1200S vibratome (Leica, Nussloch, Germany). Free-floating sections were washed in 1xPBS, blocked for 1 hr in blocking buffer (PBS containing 0.25% Triton-X-100 and 5% normal goat), then incubated with *α*-V5 primary antibody (AB3792, Millipore Sigma; 1:1000) in blocking buffer at 4°C overnight. The following day, slices were washed three times with PBST (PBS and 0.1% Tween-20), then incubated with *α*-rabbit secondary antibody (Alexa-Fluor 488 Goat *α*-Rabbit, AB150077, Thermo Fisher, 1:5000) for 2.5 hrs in blocking buffer. After this incubation, brain sections were washed three times in PBST, incubated in DAPI (Millipore, Burlington, MA; 1:2000) for 5 min, then washed three times in PBS for 5 mins immediately before mounting. Sections were mounted on Superfrost Plus Microscope slides (Fisher Scientific, Waltham, MA) using VectaShield Hard Set Mounting Medium (Vector Laboratories, Burlingame, CA). Fluorescent images were captured using a ZEISS LSM 800 confocal (Zeiss, Oberkochen, Germany). Note that immunostainings were not performed for tdTomato; instead, native fluorescence was used to visualize tdTomato in all images from *mWAKE^Cre^* mice.

### *In situ* hybridization

*In situ* hybridization was performed using the RNAscope 2.5 Chromogenic Assay and the BaseScope^TM^ Detection Reagent Kit according to manufacturer’s instructions (Advanced Cell Diagnostics/ACD, Newark, CA) (Wang et al., 2012). Briefly, WT mouse brains were dissected and fresh frozen in Tissue-Tek O.C.T. (VWR, Logan Township, NJ). Sections were then cryosectioned at 10 μm and placed onto SuperFrost Plus slides, which were then dried at −20°C for 1 hr and transferred to −80°C for storage. Prior to use, slides were immersed in pre-chilled 4% paraformaldehyde (PFA) for 15 min. Following post-fixation, slides were dried by sequential immersion in 50% EtOH, 70% EtOH, and 100% EtOH, each at room temperature for 5 min before incubation at 37°C for 30 min. Using an Immedge^TM^ hydrophobic barrier pen, boxes were drawn around each individual section and left to dry. Slides were then treated with RNAscope Hydrogen Peroxide for 10 min at room temperature and rinsed once in tap water before incubating with Protease IV for 30 min at room temperature. The slides were then washed in 1xPBS. Target probe (custom-produced by ACD to target exon 4 of *mWake*) was hybridized to tissue for 2 hr at 40°C, and seven subsequent amplification and wash steps followed, fixing peroxidase enzymes to hybridized target mRNA. Signal was detected by chromogenic reactions with the BaseScope^TM^ Fast RED, and sections were counterstained with hematoxylin. All slides from *in situ* studies were imaged on a Keyence BZ-X700 microscope (Keyence, Itasca, IL) under 10x brightfield illumination, with higher magnification images acquired via LSM 800 confocal microscope.

### scRNA-Seq data analysis

A scRNA-Seq dataset was previously generated from flow-sorted tdTomato^+^ cells of 7 week old male *mWAKE^(Cre/+)^* hypothalami (Bell et al., 2020) and used for analysis in this study (GSE146166). Data were processed as described previously (Bell et al., 2020) using Seurat V3 (Stuart et al., 2019) to perform downstream analyses. Initially, the dataset was divided into 3 distinct clusters: SCN neurons, non-SCN neurons, and ependymal cells (see Figure S3a). Non-SCN neurons were divided further based on spatial information derived from previous work (Bell et al., 2020) into 4 additional clusters: ventromedial hypothalamus, preoptic area, dorsomedial hypothalamus, and tuberomammillary nucleus.

Two major data analyses were performed in this study. First, *mWake^+^* neurons identified from our collected cells belonging to individual hypothalamic subregions were extracted and further reclustered to identify different sub-clusters of *mWake^+^* neurons. Reclustering was repeated until the child clusters could not generate any further clusters that had differential expression of neuropeptides and neurotransmitters. Second, we sought to identify *mWake^+^* neurons in existing scRNA-Seq datasets. However, due to its very low expression levels, *Ankfn1/mWake* is not faithfully detected by droplet-based scRNA-Seq methods, which have limited capture efficiency (Wang, Li, Nelson, & Nabavi, 2019). Thus, in order to identify the distribution of *mWake^+^* neurons in individual hypothalamic regions, key molecular markers of our FACS-sorted *mWake^+^* neurons were used to train existing hypothalamus scRNA-Seq datasets using Garnett (Pliner, Shendure, & Trapnell, 2019). Only scRNA-Seq datasets from male mice were used. To validate this approach, we trained a previously published VMH SMART-Seq dataset (Kim et al., 2019), where *Ankfn1/mWake* expression could be directly captured. The trained SMART-Seq dataset of putative *mWake^+^* cells was compared to *Ankfn1/mWake* gene expression and demonstrated ∼90% similarity in cell distribution (see Figure S2b,c), suggesting that the training method should efficiently identify *mWake^+^* populations in any hypothalamic dataset. The data training process was then applied to a SCN 10x Genomics and Drop-seq dataset (Wen et al., 2020) and a POA 10x Genomics dataset (Moffitt et al., 2018).

## Results

### Experimental models for investigating mWAKE expression

To perform a comprehensive examination of *mWake* expression in the murine brain, we applied three parallel strategies for visualizing *mWake* mRNA, mWAKE protein, and *mWake^+^* cells (Figure 1a-c). We previously used standard chromogenic ISH to identify *mWake* in the SCN, where its expression is relatively high (Liu et al., 2014). Here, to significantly enhance sensitivity and detect *mWake* mRNA expression more broadly, we used the RNAscope system (ACD Biosystems) (Wang et al., 2012) (Figure 1a). In order to directly visualize mWAKE protein, we made multiple attempts to generate anti-mWAKE antibodies that could be used for immunostaining, but were unsuccessful (data not shown). We and others similarly failed at producing such antibodies against the *Drosophila* WAKE protein and a related isoform (Liu et al., 2014; Mauri, Reichardt, Mummery-Widmer, Yamazaki, & Knoblich, 2014), suggesting that this family of proteins may be particularly refractory to this approach. Thus, as an alternative approach, we used CRISPR/Cas9 to generate a V5 epitope-tagged mouse (*mWAKE^V5^*), where the V5 tag is fused in-frame at the 3’ end of the mWAKE protein, immediately preceding the stop codon (Figure 1b). Lastly, to identify *mWake^+^* cells, we used a previously described *mWAKE^Cre^* mouse line (Bell et al., 2020), where exon 5 of the *mWake* gene is replaced with a cassette containing tdTomato and a split Cre-recombinase, separated by a p2A self-cleaving peptide (Figure 1c).

**Figure 1.**
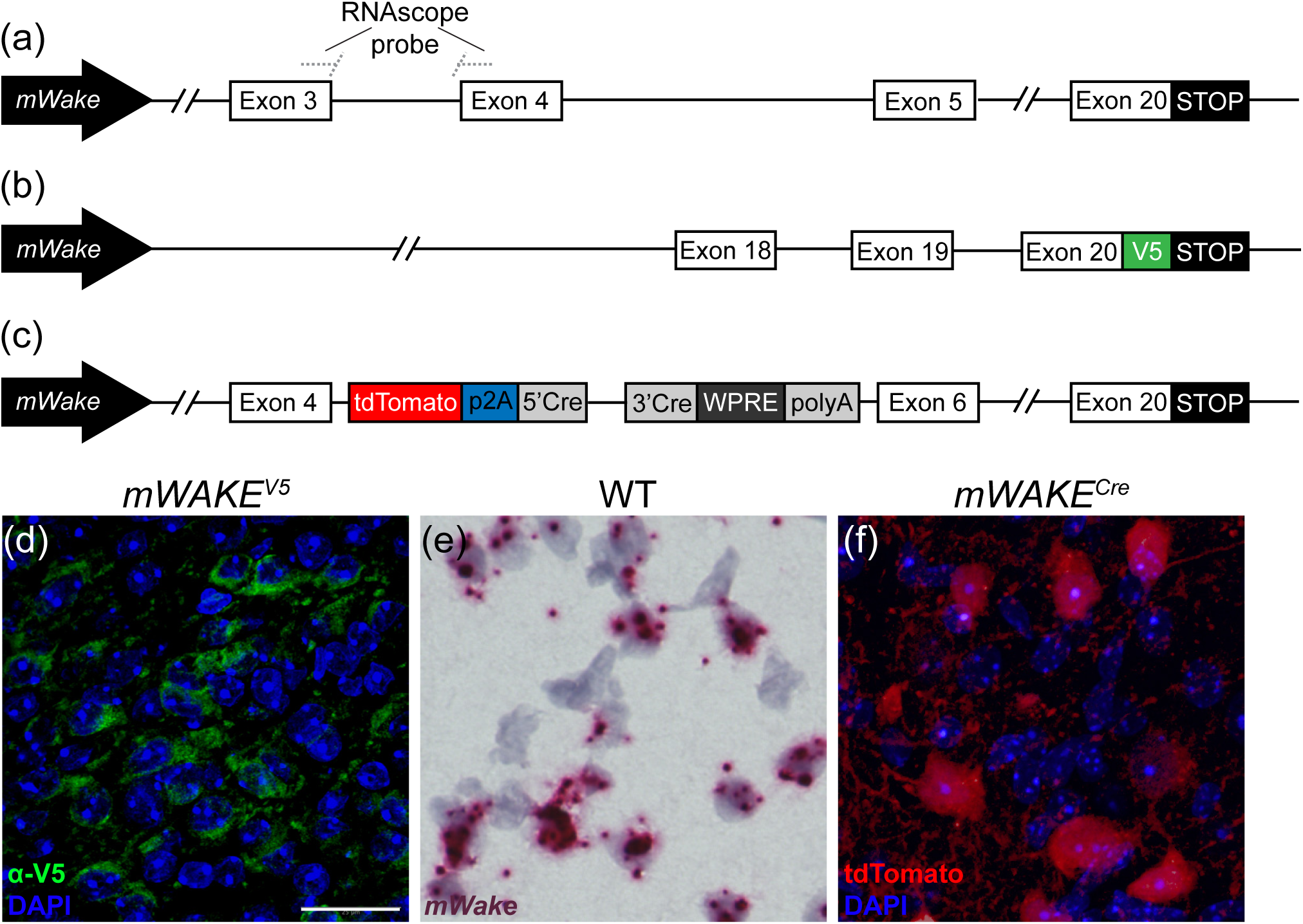
Methods for visualizing *mWake* expression. The three main approaches for characterizing *mWake* expression are shown. (a) *mWake* mRNA was visualized using RNAscope, a high sensitivity technique for ISH, using probes bridging exons 3 and 4. (b) mWAKE protein was characterized by anti-V5 immunostaining of a transgenic mouse line where an inserted V5 epitope tag was fused to the C-terminus of mWAKE. (c) *mWake^+^* cell bodies and processes were labeled by imaging tdTomato native fluorescence in a previously described transgenic mouse line (Bell et al., 2020), where exon 5 was replaced with a cassette containing both tdTomato and Cre-recombinase. Representative images from each approach are depicted. (d) IF using *α*-V5 antibodies (green) in the SCN of an *mWAKE^(V5/+)^* mouse shows mWAKE-V5 fusion protein in the cytosol. (e) Chromogenic labeling of *mWake* mRNA (red) via RNAscope ISH in a wild-type mouse. (f) SCN neurons filled with tdTomato fluorescence (red) in an *mWAKE^(Cre/+)^* mouse, labelling both the cell bodies as well as the *mWake^+^* processes. Images (d) and (f) also include the nuclear counterstain DAPI (blue), while (e) includes a hematoxylin counterstain (purple). The scale bar in (d) is 25 μm and applies to all images.

To demonstrate the fidelity of the expression patterns derived from these transgenic mouse lines, we performed 2 comparisons. First, anti-V5 staining in *mWAKE^V5^* mice and native tdTomato fluorescence in *mWAKE^(Cre/+)^* mice were both compared to *mWake* mRNA expression assessed by RNAscope at the regional level. As shown in Figures 1-14, every region exhibiting significant *mWake* expression by RNAscope also demonstrated mWAKE-V5 and tdTomato signal in *mWAKE^(V5/+)^* and *mWAKE^(Cre/+)^* mice, respectively. Second, we directly examined co-localization of mWAKE-V5 and tdTomato expression at the individual cell level in transheterozygote *mWAKE^(V5/Cre)^* mice. For the majority of regions examined, there was >90% concordance between mWAKE-V5 and tdTomato signal, with a maximum and minimum of 98% and 81% co-labeling, respectively (Figure S1). Together, these data argue that the *mWAKE^V5^* and *mWAKE^Cre^* mice serve as valuable models for investigating the expression pattern of mWAKE in the murine brain.

Each approach provides specific and complementary information about *mWake* and *mWake*^+^ cells. mWAKE-V5 allows for visualization of the protein, and mWAKE-V5 signal is primarily found in the cytosol, indicating that mWAKE protein is neither confined to the nucleus nor predominantly trafficked to synaptic terminals (Figure 1d). Not only does RNAscope directly label *mWake* mRNA, but because RNAscope signal is strongest in the nucleus in most cells, it is helpful for visualizing m*Wake^+^* cell bodies in projection-dense regions (Figure 1e). Finally, tdTomato in *mWAKE^Cre^* mice fills the cytosol of putative *mWake^+^* cells, and is therefore well-suited for visualizing projections of *mWake^+^* neurons (Figure 1f). Below, for all of the *mWake^+^* regions identified across the brain, we utilized these 3 approaches to characterize *mWake* expression and *mWake^+^* cells. Because of the substantial *mWake* expression in the hypothalamus, we discuss this region first, followed by other subcortical regions, the brainstem, and the cortex. Finally, we describe *mWake* expression in areas of the brain that, to the best of our knowledge, have not been previously named.

### Characterization of *mWake*^+^ populations in the hypothalamus

#### Overview

The hypothalamus is a subcortical region involved in the homeostatic and circadian regulation of wide array of physiological processes and behaviors (McKinsey, Ahmed, & Shah, 2018; Saper, Lu, Chou, & Gooley, 2005; Sisley & Sandoval, 2011; Tan & Knight, 2018). *mWake* exhibits significant expression in multiple regions of the hypothalamus, with notable enrichment in the SCN, the circadian pacemaker in mammals (Bell et al., 2020; Liu et al., 2014; Zhang et al., 2015). Beyond the 3 main approaches outlined above, we also analyzed scRNA-Seq data from *mWake^+^* cells in the hypothalamus to gain deeper insights into the molecular identities of these cells. We previously generated a scRNA-Seq dataset from tdTomato-labeled cells sorted from dissected hypothalami of *mWAKE^(Cre/+)^* mice (Bell et al., 2020), and here we perform additional analyses to further discriminate the identity of *mWake*^+^ populations. Specifically, we performed location-specific reclustering to sub-divide *mWake^+^* cells within a given hypothalamic region and also identified *mWake^+^* neuronal subsets within previously published scRNA-Seq datasets.

Since single-cell expression profiles in published scRNA-Seq datasets often do not capture *mWake* due to its low endogenous expression (Zhang et al., 2015) (see Methods), we derived and used key molecular markers for *mWake^+^* hypothalamic neurons to identify putative *mWake^+^* cells in these datasets. Specifically, we are aware of only one published scRNA-Seq dataset from hypothalamic tissue (which used the highly sensitive SMART-Seq protocol on VMH tissue (Kim et al., 2019)) where *Ankfn1/mWake* expression is captured, and thus *mWake^+^* cells can be directly identified. Using the expression profiles of our FACS-sorted *mWake^+^* cells, we generated a list of key genetic markers that putatively define *mWake*-expressing cells without the need for *mWake* mRNA capture. We found a strong correlation (89.9%) between the transcriptional profiles of the *Ankfn1/mWake-*expressing cells from Kim et al. (2019) and those from their dataset we identified as *mWake^+^* using our predictive genetic markers (Figure S2b,c). This finding validates our approach and argues that we should be able to reliably identify *mWake^+^* cells within other published scRNA-Seq datasets.

The tSNE plot containing all sequenced *mWake^+^* cells from our FACS-sorted tdTomato^+^ dataset reveals three distinct *mWake^+^* clusters containing SCN neuron, non-SCN neuron, and non-neuronal ependymal cell populations (Figure S3a). The *mWake*^+^ SCN neuron cluster demonstrates marked enrichment of the neuropeptides vasoactive intestinal peptide (*Vip*) and gastrin-releasing peptide (*Grp*), as well as the transcription factors *Lhx1* and *Six3* (Bedont et al., 2017; Bell et al., 2020; Welsh, Takahashi, & Kay, 2010). The non-SCN mWAKE neurons are a heterogenous population containing both GABAergic or glutamatergic populations (Bell et al., 2020). As discussed below, these non-SCN *mWake*^+^ neurons are found in the dorsomedial hypothalamus (DMH), ventromedial hypothalamus (VMH), POA, tuberomammillary nucleus (TMN), and lateral hypothalamus (LH). Finally, the *mWake^+^* ependymal cell group exhibits substantial expression of the ependymal cell markers forkhead box J1 (*Foxj1*) and retinoic acid receptor responder 2 (*Rarres2*) (Figure S3b) (Miranda-Angulo, Byerly, Mesa, Wang, & Blackshaw, 2014; Shah et al., 2018).

#### Suprachiasmatic nucleus

We and others previously demonstrated that *mWake* mRNA expression is enriched in the SCN (Bell et al., 2020; Liu et al., 2014; Zhang et al., 2015). The SCN can be roughly divided into a ventral “core” (SCNc) region and a dorsal “shell” (SCNs) region, where the former receives direct projections from intrinsically-photosensitive retinal ganglion cells (ipRGC) in the retina via the retinohypothalamic tract (Altimus et al., 2008; Welsh et al., 2010; Yan et al., 2007). Staining for the mWAKE-V5 fusion protein reveals that a majority of the central core neurons of the SCN express mWAKE protein, and endogenous tdTomato fluorescence indicates these cell bodies are intermingled with axons and processes stemming from *mWake^+^* neurons (Figure 2a-c). The density of these cell bodies is highest in the ventromedial limit of the core, but there are additional cells with V5 signal and nuclei positive for *mWake* mRNA around the outermost dorsal perimeter of the core. Additionally, the dorsal apex of the shell contains a population of tdTomato^+^ fibers, which likely stem from *mWake^+^* neurons in the core, although we cannot rule out other *mWake^+^* cells projecting to this region (Figure 2c).

**Figure 2.**
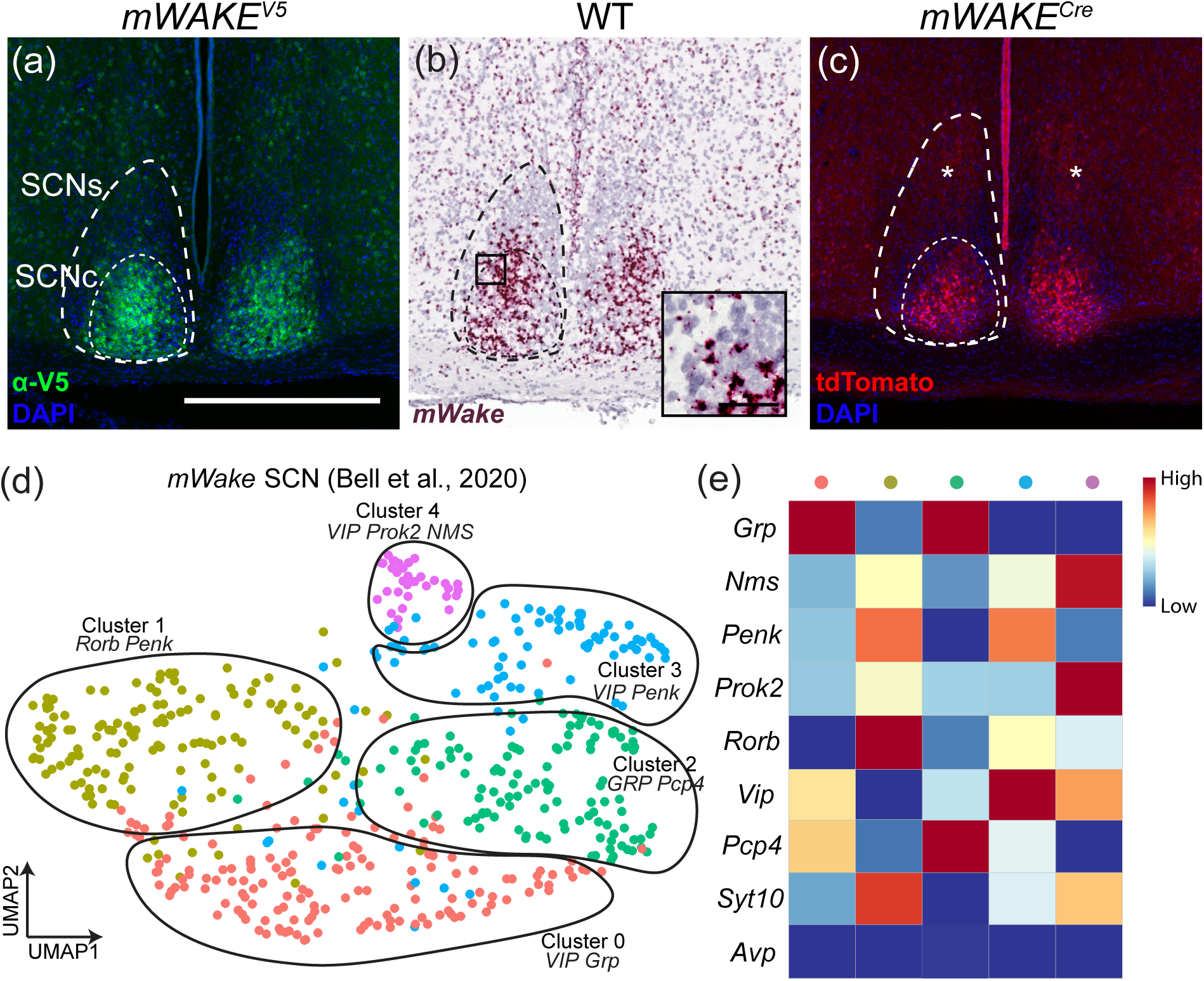
Visualization and subclustering of *mWake^+^* neurons in the SCN. Representative images of *mWake* expression in the SCN are shown. (a) Anti-V5 IF staining (green) with DAPI in *mWAKE^(V5/+)^* mice. (b) Chromogenic RNAscope ISH for *mWake* mRNA (red) with hematoxylin counterstain (purple) in wild-type (WT) mice. (c) Endogenous tdTomato fluorescence (red) in *mWAKE^(Cre/+)^* mice with DAPI. In (a)-(c), the larger dashed line denotes the extent of the SCN shell (SCNs), while the thinner inner dashed line outlines the SCN core (SCNc). The black box in (b) depicts the approximate location of the representative higher-magnification inset in the lower right corner. The asterisks in (c) indicate the tdTomato^+^ processes in the dorsal SCN shell. The scale bar in (a) is equivalent to 500 μm, and applies across the row, and the scale bar in the inset of (b) denotes 50 μm. (d) UMAP plot of *mWake^SCN^* neurons reveals five subgroups with key gene markers: cluster 0 (*Vip, Grp*, red); cluster 1 (*Rorb, Penk*, yellow); cluster 2 (*Grp, Pcp4*, green); cluster 3 (*Vip, Penk*, blue); cluster 4 (*Vip, Prok2, Nms*, purple). (e) Heatmap showing expression levels for selected molecular markers of the 5 clusters shown in (d).

Next, we examined the expression profiles of the *mWake^+^* SCN neurons identified from our scRNA-Seq dataset. We previously identified multiple subpopulations of *mWake^+^* SCN neurons directly from the entire hypothalamic population (Bell et al., 2020). Here, we first isolated *mWake^+^* SCN neurons and then further stratified this population to reveal 5 main clusters identified by distinct expression patterns of SCN neuropeptides (Figure 2d). *mWake^SCN^* neurons are generally enriched for neuropeptide markers of the SCNc (e.g., *Vip* and *Grp*), but do not significantly express the primary SCNs marker arginine vasopressin *Avp* (Abrahamson & Moore, 2001; Welsh et al., 2010; Yan et al., 2007) (Figure 2d,e). Two *mWake^SCN^* clusters (clusters 3 and 4) express moderate to high levels of *Vip*, with little *Grp* expression. These two clusters likely represent the most ventral subgroups, given that *Vip* expression marks ventral SCN neurons. *Grp* labels ventromedial SCN neurons in the core region and is highly expressed by two *mWake^SCN^* clusters (clusters 0 and 2), one of which also demonstrates moderate expression of *Vip* (cluster 0). Finally, one cluster (cluster 1) does not exhibit significant levels of either *Vip* or *Grp*, but instead is enriched for pro-enkephalin (*Penk*) and retinoic acid-related orphan receptor *β* (*Rorb*), suggesting a more dorsal location within the SCNc (Wen et al., 2020). Additional SCN markers include the following: neuromedin S (*Nms*) (Lee et al., 2015), which is expressed at moderate to high levels in clusters 1, 3, and 4; prokineticin 2 (*Prok2*) (Cheng et al., 2002), which is expressed at moderate to high levels in clusters 1 and 4; and synaptotagmin 10 (*Syt10*) (Husse, Zhou, Shostak, Oster, & Eichele, 2011), which is expressed at moderate to high levels in clusters 1, 3 and 4.

In order to distinguish *mWake^SCN^* cells within the entire population of SCN neurons, we identified putative *mWake^+^* cells from a published whole-SCN scRNA-Seq dataset and reclustered their data (Figure S4) (Wen et al., 2020). The cells from this population which express the key molecular markers for *mWake* primarily occupy those clusters which demarcate SCN core neurons, including ∼90% of cells in *Vip*^+^ and *Grp*^+^ clusters, and ∼70% of cells in the clusters expressing *Penk*. These analyses underscore the diversity of *mWake^SCN^* neuron identity within an anatomically restricted population. Moreover, given the proposed role of the SCNc in light-dependent circadian processes (Herzog, Hermanstyne, Smyllie, & Hastings, 2017; Yan et al., 2007), these findings suggest that mWAKE regulates light-dependent behaviors.

#### Dorsomedial hypothalamus

Outside of the SCN, we also observed *mWake^+^* cells throughout the hypothalamus using our mouse models and RNAscope ISH, as well as from spatial information encoded in scRNA-Seq transcript data (Bell et al., 2020). The DMH plays a key role in circadian-dependent arousal and output behaviors, including sleep-wake patterns, locomotor activity, and the timing of feeding (Saper, Lu, Chou, & Gooley, 2005). *mWake^+^* neurons within this nucleus exhibit greater spiking frequency during the night compared to the day and promote arousal, likely via outputs to noradrenergic locus coeruleus (LC) neurons and processes broadly projecting throughout the mouse brain (Bell et al., 2020). *mWake* ISH and staining for V5 reveals cell bodies distributed throughout much of the DMH, with increased density in the ventral portion (Figure 3a,b). tdTomato-expressing processes are not significantly enriched within the DMH region compared to the rest of the hypothalamus (Figure 3c). We previously demonstrated the broad projection patterns of these neurons, targeting regions including the BF, striatum, brainstem, and the corpus callosum (cc) white matter tract underlying the neocortex (Bell et al., 2020).

**Figure 3.**
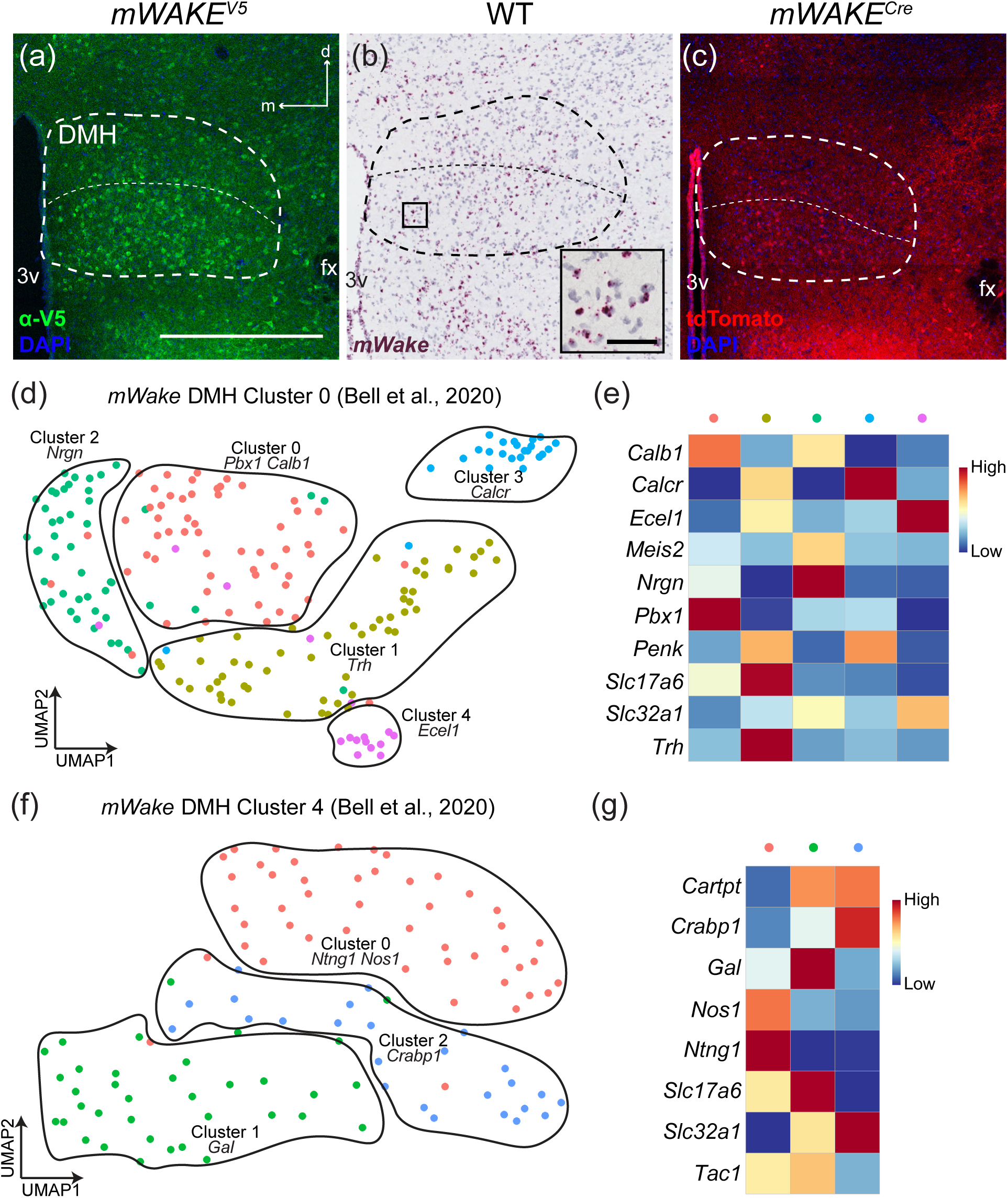
*mWake^+^* cells and scRNAseq populations in the DMH. Representative images of *mWake* expression in the DMH are shown. (a) anti-V5 IF staining (green) with DAPI in *mWAKE^(V5/+)^* mice. (b) Chromogenic RNAscope ISH for *mWake* mRNA (red) with hematoxylin counterstain (purple) in wild-type (WT) mice. (c) Endogenous tdTomato fluorescence (red) in *mWAKE^(Cre/+)^* mice with DAPI. In (a)-(c), the large dashed outline denotes the boundary of the DMH, with a thinner dashed horizontal line dividing the dorsal from the ventral portion. The black box in (b) depicts the approximate location of the representative higher-magnification inset in the lower right corner. In (a), the scale bar is equivalent to 500 μm, and the arrows indicate dorsal (d) and medial (m) directions, and both also apply to (b) and (c). The scale bar in the inset of (b) denotes 50 μm. (d) and (f) UMAP plots of two *mWake^DMH^* populations, with 5 and 3 subclusters, respectively. (e) and (g) Heatmaps showing expression levels for key molecular markers for the subclusters shown in (d) and (f), respectively. For (d) and (e), the original population was named ‘Cluster 0’ in Bell *et. al* (2020), and the 5 subclusters are defined as cluster 0 (*Pbx, Calb1*, red), cluster 1 (*Trh*, yellow), cluster 2 (*Nrgn*, green), cluster 3 (*Calcr*, blue), cluster 4 (*Ecel1*, purple). For (f) and (g), the original population was called ‘Cluster 4’ in Bell *et. al* (2020), and the 3 subclusters are defined as cluster 0 (*Ntng1, Nos1*, red), cluster 1 (*Gal*, green), cluster 2 (*Crabp1*, blue). Abbreviations: DMH, dorsomedial hypothalamus; fx, fornix; 3v, third ventricle.

Our original scRNA-Seq clustering revealed two main subpopulations of *mWake^DMH^* neurons, which were primarily identified as glutamatergic and expressing a similar constellation of genes, including cholecystokinin (*Cck*) and prepronociceptin (*Pnoc*) (Bell et al., 2020). Here, we have refined our classification of *mWake^DMH^* neurons (Figure 3d-g) and further separated these two clusters. The original *mWake^DMH^* cluster 0 neurons can be sub-divided into five sub-clusters: two are strongly glutamatergic (express *Slc17a6/Vglut2*), one of which strongly expresses Thyrotropin releasing hormone (*Trh*); two are GABAergic (express *Slc32A1/Vgat*); and one cluster which is not clearly glutamatergic or GABAergic, but does express *Penk*. Further stratification of *mWake^DMH^* cluster 4 neurons into three groups reveals one GABAergic subcluster, one with expression of both glutamatergic and GABAergic markers as well as galanin (*Gal*), and one that is glutamatergic and enriched for nitric oxide synthase 1 (*Nos1*), proposed as a marker of hypothalamic neurons involved in regulation of energy homeostasis (Rupp et al., 2018).

#### Ventromedial hypothalamus

The VMH has been implicated in the regulation of a variety of motivated behaviors, including appetitive-drive, aggression, and sexual function (Hashikawa, Hashikawa, Falkner, & Lin, 2017; King, 2006; Yang et al., 2013). *mWake^VMH^* cell bodies, labelled by V5 fusion protein and particularly ISH, are largely constrained to the ventrolateral portion of the region, which has previously been implicated in circadian regulation of aggression (Todd et al., 2018). In contrast, nearly the entire VMH is densely covered by tdTomato^+^ projections (Figure 4a-c).

**Figure 4.**
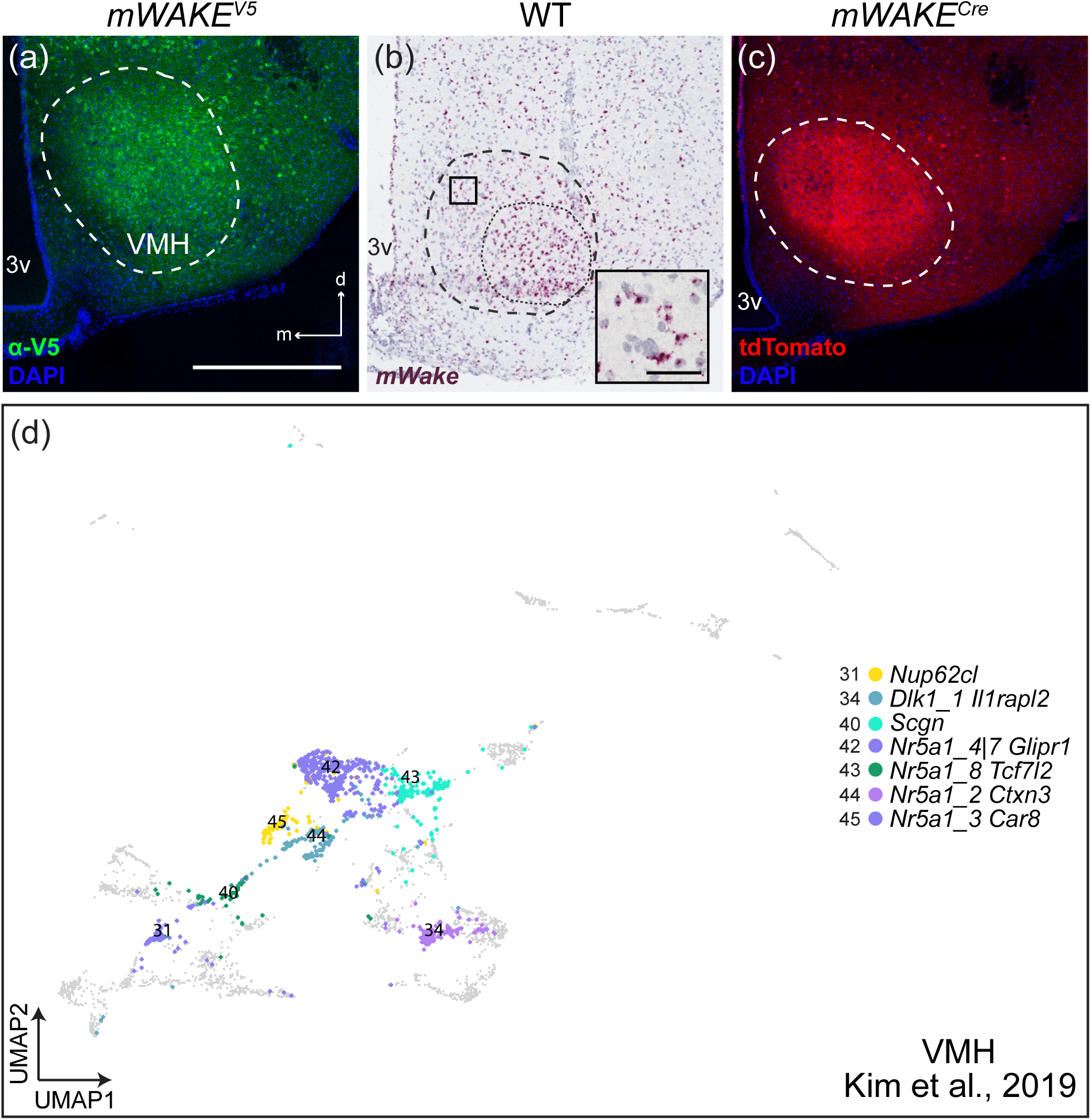
*mWake^+^* cells and scRNAseq populations in the VMH. Representative images of *mWake* expression in the VMH are shown. (a) Anti-V5 IF staining (green) with DAPI in *mWAKE^(V5/+)^* mice. (b) Chromogenic RNAscope ISH for *mWake* mRNA (red) with hematoxylin counterstain (purple) in wild-type (WT) mice. (c) Endogenous tdTomato fluorescence (red) in *mWAKE^(Cre/+)^* mice with DAPI. In (a)-(c), the larger dashed line denotes the boundary of the VMH and in (b) a thinner dashed line encircles the ventrolateral region. In (a), the scale bar is equivalent to 500 μm, and the arrows indicate the dorsal (d) and medial (m) directions, and both also apply to (b) and (c). The scale bar in the inset of (b) denotes 50 μm. (d) UMAP plot of all cells from Kim *et. al.* (2019) re-clustered. Clusters where putative *mWake*^+^ cells constitute >70% of the population are colored and represent 7 out of 46 identified transcriptional profiles: cluster 31 (*Nup62cl*, yellow), cluster 34 (*Dlk1, Il1rapl2*, steel blue), cluster 40 (*Scgn*, teal blue), cluster 42 (*Nr5a1, Glipr1*, dark purple), cluster 43 (*Nr5a1, Tcf7l2*, green), cluster 44 (*Nr5a1, Ctxn3*, light purple), cluster 45 (*Nr5a1, Car8*, deep purple). Abbreviations: 3v, third ventricle; VMH, ventromedial hypothalamus.

While the low abundance of *mWake^+^* neurons in the VMH prevented further subclustering of our previously defined scRNA-Seq VMH cluster, we identified *mWake^+^* cells from a previously published VMH scRNA-Seq dataset (Kim et al., 2019) as described above. Kim et al. (2019) performed a systematic analysis of VMH neurons using 10x Genomics and SMART-seq scRNA-Seq, combined with seq-FISH, and classified VMH neurons into 46 distinct clusters. Putative *mWake^+^* neurons were broadly distributed throughout these clusters, but 7 of these clusters contained high (>70%) numbers of these neurons (Figure 4d). Of these, four groups were primarily defined by their expression of nuclear receptor subfamily 5 group a member 1 (*Nr5a1*), and one by secretagogin (*Scgn*); these groups were largely constrained to the central VMH (Kim et al., 2019). Two additional clusters with high percentages of *mWake^+^* neurons are defined by expression of Nucleoporin-62 C-terminal-like (*Nup62cl*) and Delta-like 1 homolog (*Dlk1*), and both primarily consisted of cells from the ventrolateral VMH. Interestingly, despite extensive testing, these individual transcriptionally-defined VMH clusters did not exhibit distinct projection patterns or specific roles in behavior (Kim et al., 2019).

#### Pre-optic area

The POA occupies the transitional edge of the hypothalamus where its rostral terminus merges with structures of the BF. Circuits in the POA region are involved in thermoregulation, sexually-dimorphic reproductive behaviors, and control of sleep-wake states (Gaus, Strecker, Tate, Parker, & Saper, 2002; Wei et al., 2018; Zhao et al., 2017). Interestingly, we observed no cell bodies (*mWake* mRNA or V5 staining) and no tdTomato^+^ processes in the ventrolateral preoptic area (VLPO), a well-established sleep regulatory center. Instead, cell bodies and processes are found where the BF and hypothalamus meet, with a particularly dense ventromedial grouping immediately rostral to the SCN, nearly lining the ventral portion of the 3v (Figure 5a,b). In addition to these cell bodies, there appears to be a lateral band of tdTomato^+^ projections immediately at the interface of the gray matter of the POA and the white matter tract forming the optic chiasm (oc) below (Figure 5c).

**Figure 5.**
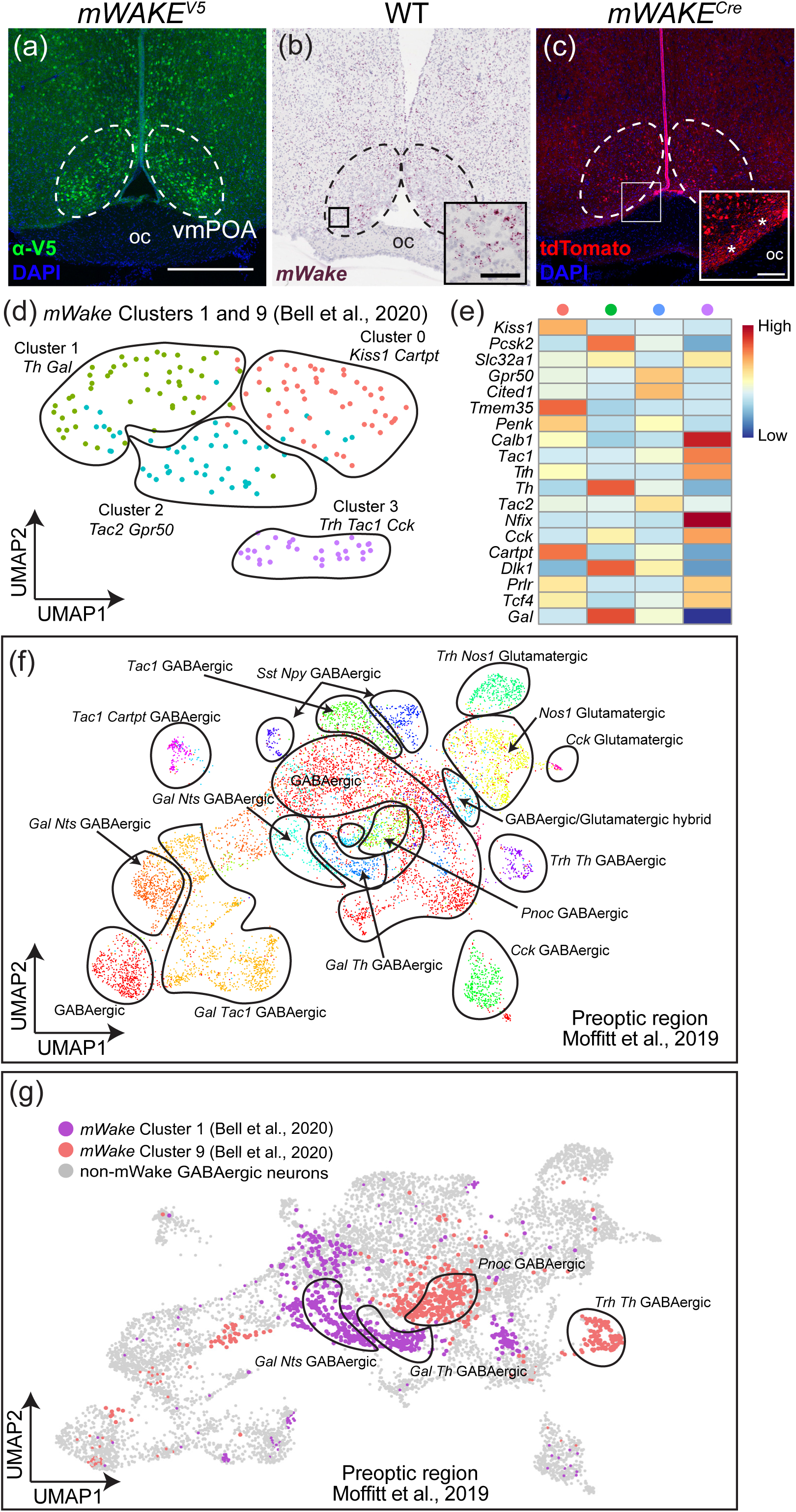
*mWake^+^* cells and scRNAseq populations in the POA. Representative images of *mWake* expression in the POA are shown. (a) anti-V5 IF staining (green) with DAPI in *mWAKE^(V5/+)^* mice. (b) Chromogenic RNAscope ISH for *mWake* mRNA (red) with hematoxylin counterstain (purple) in wild-type (WT) mice. (c) endogenous tdTomato fluorescence (red) in *mWAKE^(Cre/+)^* mice with DAPI. In (a)-(c), the dashed outlines denote the ventromedial POA. The black box in (b) and the white box in (c) depict the approximate location of the representative higher-magnification insets in the lower right corner of each. In the inset of (c), asterisks indicate the tdTomato^+^ processes. The scale bar in (a) is equivalent to 500 μm and applies to (b) and (c). Scale bars for the insets in (b) and (c) represent 50 μm and 100 μm, respectively. d) UMAP plot of *mWake^POA^* neurons demonstrating 4 sub-clusters marked by expression of key genes: cluster 0 (*Kiss1, Cartpt*, red), cluster 1 (*Th, Gal*, yellow), cluster 2 (*Tac2, Gpr50*, blue), cluster 3 (*Trh, Tac1, Cck*, purple). (e) Heatmap showing expression levels for select molecular markers for the clusters shown in (d). (f) UMAP plot of all POA neurons from Moffit *et. al.* (2019), with key genetic markers and glutamatergic/GABAergic identity indicated for each. (g) UMAP plot of all POA neurons from Moffit *et. al.* (2019), where 4 clusters with >90% putative *mWake^+^* cells highlighted in color. These clusters correspond to ‘cluster 1’ (purple) and ‘cluster 9’ (red) from Bell *et. al.* (2020), are all GABAergic, and defined by the following markers: *Gal* and *Nts*; *Gal* and *Th*; *Trh* and *Th*; and *Pnoc*. Abbreviations: oc, optic chiasm; vmPOA, ventromedial preoptic area.

Our scRNA-Seq reclustering analysis takes the two original POA groups (clusters 1 and 9 (Bell et al., 2020)), and delineates four subgroups of *mWake^POA^* neurons based on expression patterns of POA-relevant genes (Figure 5d,e). Three contain primarily GABAergic *mWake^+^* neurons and two express *Gal*. Classically, *Gal^+^* neurons in the POA were thought to be sleep-promoting and located in VLPO, but recently an additional *Gal*-expressing population in the POA has been shown to stimulate wakefulness, a circuit which may include *mWake^+^* cells (Chung et al., 2017; Kroeger et al., 2018). We next applied our training algorithm to a published scRNA-Seq dataset of all POA neurons (Moffitt et al., 2018). *mWake* expression is largely localized to four clusters within their population (where *mWake^+^* cells constitute >90% cells) and is found in <10% of cells in the other clusters (Figure 5f,g). All of these groups were GABAergic, with two co-expressing *Gal*, and the other two expressing *Trh* or *Pnoc*. Interestingly, one of our clusters, and two clusters of putative *mWake^POA^* cells, express tyrosine hydroxylase (*TH*), raising the possibility that *mWake* may label dopaminergic or noradrenergic neurons.

#### Tuberomammillary nucleus

The TMN contains nearly all of the histaminergic neurons in the mammalian brain, and their widespread projections have been implicated in the maintenance of waking states and attentional vigilance (Fujita et al., 2017; Gerashchenko, Chou, Blanco-Centurion, Saper, & Shiromani, 2004). *mWake^+^* neurons intermingle with, but are distinct from, this neuronal population (Bell et al., 2020). *mWAKE*-expressing cell bodies are found in two sub-regions of the TMN, one which lines the ventral-most portion of the lateral hypothalamus underneath the fornix (fx) (Figure 6a-c), and a medial group adjacent to the third ventricle (3v) above the arcuate nucleus (Figure 6d-f); both of these regions generally contain histaminergic cell bodies. The highest density of cell bodies as seen by *mWake^+^* ISH signal and V5 staining are in the medial groups adjacent to the 3v, but the tdTomato^+^ projections appear densest in the perifornicular area, delineating the boundaries of the white matter tract (Figure 6c).

**Figure 6.**
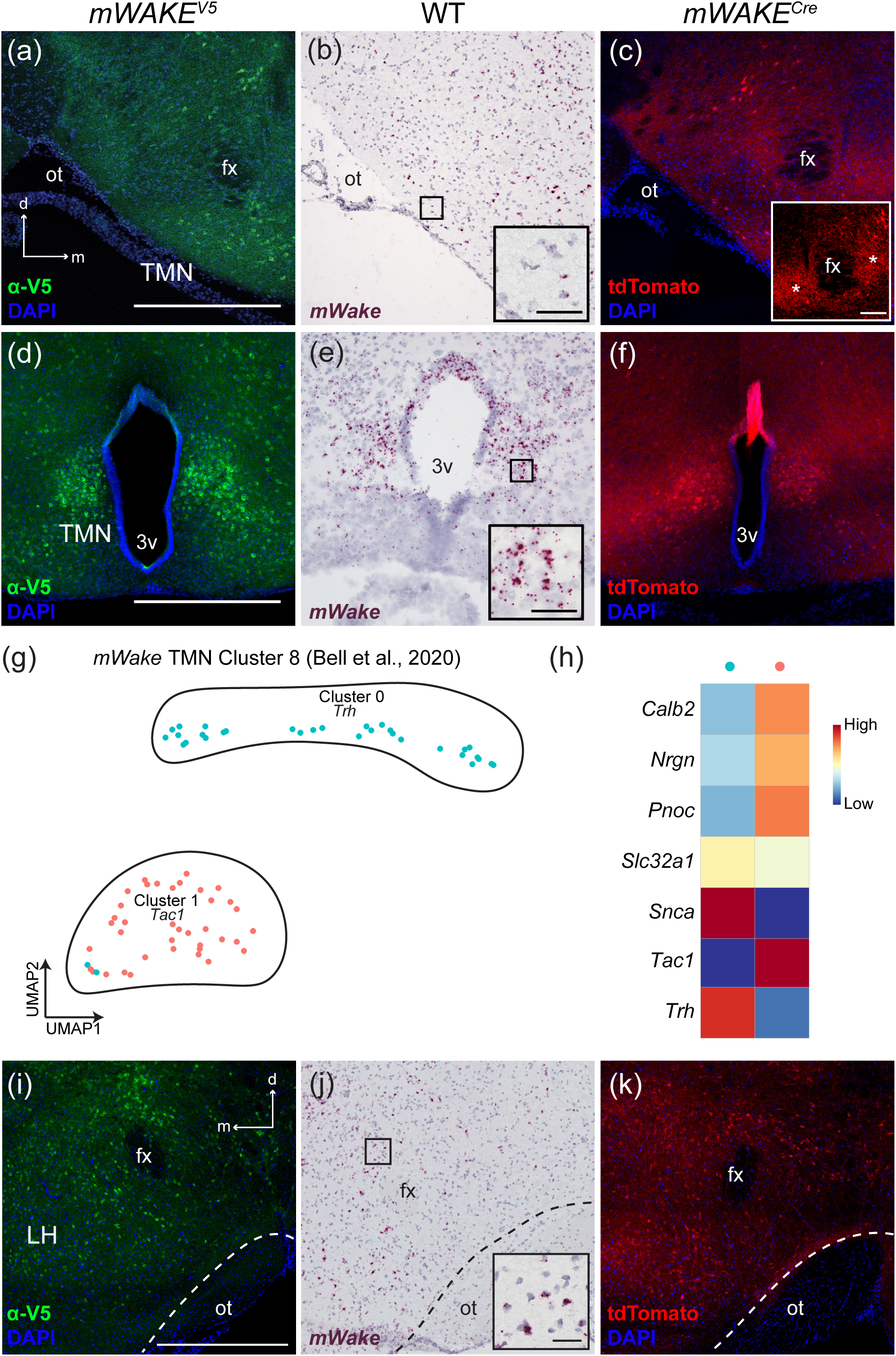
*mWake^+^* cells in the TMN and LH. Representative images of *mWake* expression in 2 areas of the TMN are shown (the perifornicular region and the medial 3v region). (a) and (d) anti-V5 IF staining (green) with DAPI in *mWAKE^(V5/+)^* mice. (b) and (e) chromogenic RNAscope ISH for *mWake* mRNA (red) with hematoxylin counterstain (purple) in wild-type (WT) mice. (c) and (f) endogenous tdTomato fluorescence (red) in *mWAKE^(Cre/+)^* mice with DAPI. Scale bars in main panels denote 500 μm and apply to other main panels across a row. The arrows in (a) indicate the dorsal (d) and medial (m) directions and also apply to (b) and (c). The black boxes in (b) and (e) depict the approximate location of the representative higher-magnification insets in the lower right corner of each. In (c), the inset shows a higher-contrast image with the tdTomato^+^ processes marked with asterisks. Scale bars in the insets of (b) and (e) denote 50 μm, and 100 μm for the inset in (c). (g) UMAP plot of *mWake^TMN^* neurons depicting 2 sub-clusters and their key genetic markers: cluster 0 (*Trh*, blue), cluster 1 (*Tac1*, red). (h) Heatmap showing expression levels for select molecular markers in the 2 clusters shown in (g). (i)-(k) Representative images of *mWake* expression in the LH area, with descriptions corresponding to (a)-(c). Dashed lines in (i)-(k) demarcate the edge of the optic tract along the ventral edge of the hypothalamus. The black box in (j) depicts the approximate location of the representative higher-magnification inset in the lower right corner, where the scale bar denotes 50 μm. Abbreviations: 3v, third ventricle; fx, fornix; LH, lateral hypothalamus; ot, optic tract; TMN, tuberomammillary nucleus.

The anatomical division of *mWake^TMN^* neurons into two groups appears to be recapitulated by our scRNA-Seq clustering analysis. This evaluation yields only two sub-clusters of *mWake^TMN^* neurons which both express *VGlut2*, but which can be distinguished by their expression of *Trh* (cluster 0) or *Pnoc* and *Tac1* (cluster 1) (Figure 6g,h). To characterize the relationship between these clusters and the two anatomical subsets, we used the Allen Brain Atlas ISH database to locate expression of key genes from each cluster within the TMN. *Trh* expression is densest along the 3v, while *Tac1* expression is primarily more ventral and lateral. These data suggest that the scRNA-Seq *mWake^TMN^* cluster 0 subgroup corresponds to the anatomically-defined 3v-adjacent medial group (Figure 6d-f), while the cluster 1 subgroup encompasses the more ventral portion of the TMN (Figure 6a-c). Currently, there are no known functional differences between these anatomically and genetically distinct regions of the TMN.

#### Lateral hypothalamus

We observed sparsely distributed *mWake^+^* cell bodies, labeled by anti-V5 and RNAscope ISH areas with dense tdTomato^+^ processes, throughout the LH (Figures 6i-k). In particular, the dorsolateral quadrant of the LH contains dense *mWake^+^* processes, as well as a population of cell bodies which are intermingled with the orexinergic cell bodies which classically occupy this region (Bell et al., 2020). The LH contains a variety of arousal-promoting circuits and cell types in addition to orexin, including wake-promoting populations of both glutamatergic and GABAergic neurons (Heiss, Yamanaka, & Kilduff, 2018; Herrera et al., 2016; Tyree, Borniger, & de Lecea, 2018; Venner, Anaclet, Broadhurst, Saper, & Fuller, 2016).

### Subcortical areas expressing mWAKE

#### Zona incerta

Outside of the hypothalamus, there are a number of subcortical regions that contain *mWake^+^* cell-bodies and processes, including the zona incerta (ZI), the BF, the caudate putamen nucleus (CPu), the subparafascicular nucleus (SPFp), the subfornical organ (SFO), and multiple regions within the limbic system. Immediately dorsal to the hypothalamus, *mWake^+^* neurons and processes label the ZI, a subthalamic gray matter tract implicated in integration of multisensory inputs, regulation of behavioral outputs, and modulation of plasticity (Figure 7a-c) (Wang, Chou, Zhang, & Tao, 2020). We observed V5 staining and *mWake* mRNA in cell bodies throughout this tract, as well as tdTomato-fluorescent projections stretching lengthwise across the area. The destinations of projections from *mWake^ZI^* neurons remain unclear, but many ZI subregions are defined by their projection pattern and/or role in behavior, such as the sleep-promoting *Lhx6^+^* cells, which extend GABAergic projections to inhibit orexinergic and GABAergic neurons in the LH, and more distant arousal-promoting neuronal subtypes in the midbrain (Liu et al., 2017; Mitrofanis, 2005).

**Figure 7.**
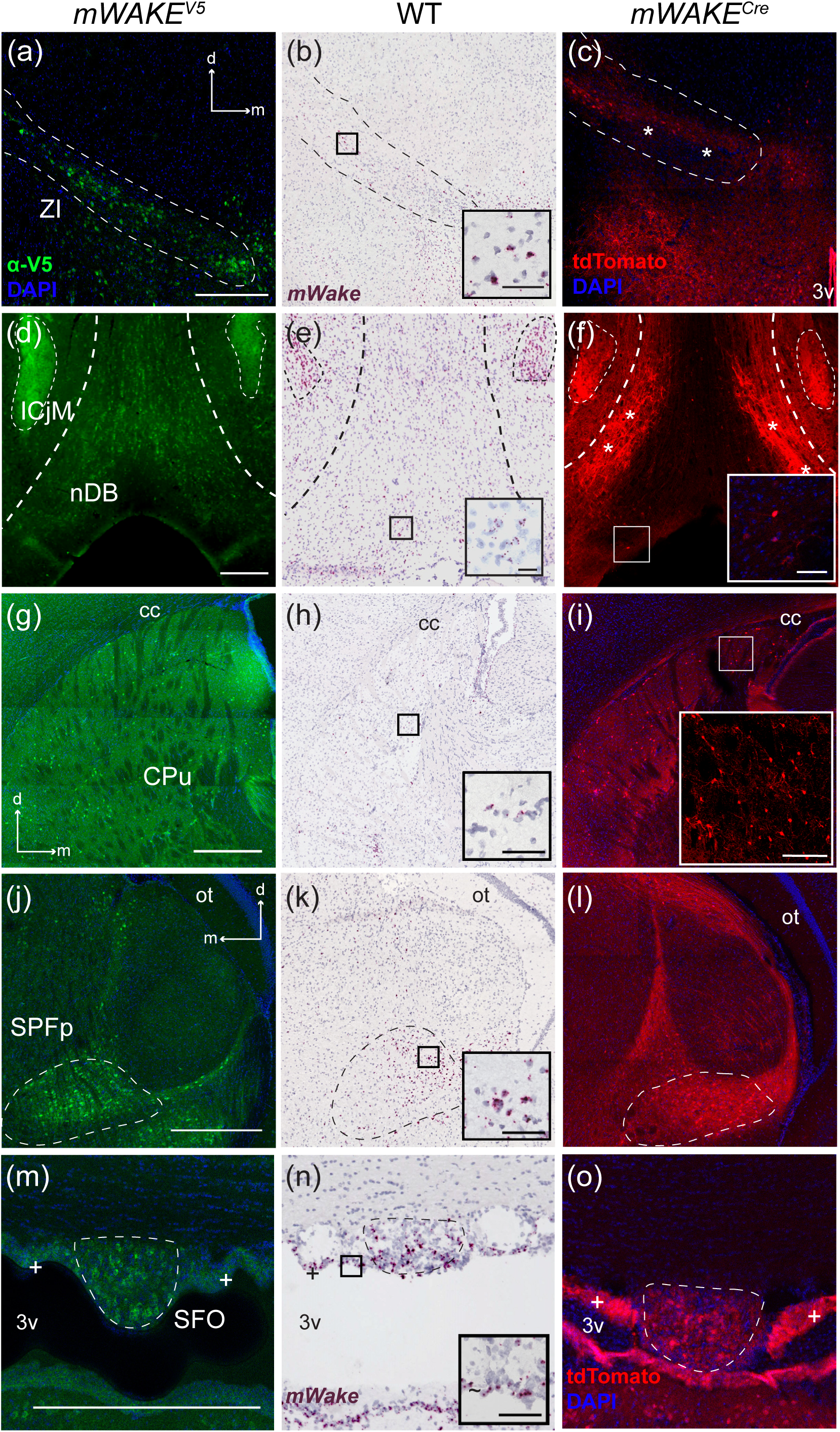
*mWake* expression in subcortical regions. Representative images of *mWake* expression in subcortical areas. Images of a ZI (a-c), the nDB (c-f), a CPu (g-i), a SPFp (j-l), and the SFO (m-o) are shown. In (a-c), the dashed line outlines the ZI. In (d-f), thicker dashed lines mark the perimeter of the nDB, and the thinner dashed line encircles the ICjM. Dashed lines outline the SPFp (j-l) and SFO (m-o), respectively. Asterisks label tdTomato^+^ processes in (c) and (f). “+” symbols in (m-o) indicate *mWake^+^* ependymal cells. (a), (d), (g), (j), and (m) Anti-V5 IF staining (green) with DAPI in *mWAKE^(V5/+)^* mice. (b), (e), (h), (k), and (n) Chromogenic RNAscope ISH for *mWake* mRNA (red) with hematoxylin counterstain (purple) in wild-type (WT) mice. (c), (f), (i), (l), and (o) Endogenous tdTomato fluorescence (red) in *mWAKE^(Cre/+)^* mice with DAPI. The black boxes in (b), (e), (h), (k), and (n) depict the approximate location of the representative higher-magnification inset in the lower right corner, where the scale bar denotes 50 μm. White boxes in (f) and (i) indicate the approximate location of the representative higher-magnification inset in the lower right corner, where the scale bars denote 50 μm and 100 μm, respectively. Scale bars in (a), (d), (g), (j), and (m) denote 500 μm and apply to other main panels across a row. Arrows in (a), (g), and (j) indicate the dorsal (d) and medial (m) directions, and apply to (b) and (c), (h) and (i), and (k) and (l), respectively. Abbreviations: 3v, third ventricle; cc, corpus collosum; CPu, caudate putamen; ICjM, Islands of Calleja insula magna; nDB, nucleus of the diagonal band; ot, optic tract; SFO, subfornicular organ; SPFp, subparafascicular nucleus parvicellular part; ZI, zona incerta.

#### Basal forebrain

The BF comprises a collection of subcortical regions arranged within the generally ventromedial space rostral to the hypothalamus, including the ventral pallidum, nucleus of the diagonal band (nDB), and substantia innominata. These regions contain a mixture of glutamatergic, GABAergic, and cholinergic neurons, which are all involved in maintenance of arousal and cortical synchrony (Anaclet et al., 2015; Kim et al., 2015; Xu et al., 2015). Within the BF, ISH and V5 staining labeled cell bodies distributed along the ventral portion of the nDB extending into the horizontal limbs, but not upwards through the vertical limb (Figure 7d-f). Additionally, tdTomato fluorescence revealed dense projections from *mWake^+^* neurons filling the lateral aspects of the nDB where there are few cell bodies (Figure 7f). Our previous data showed that *mWake^BF^* neurons are not cholinergic (Bell et al., 2020), suggesting that they may belong to the local glutamatergic and/or GABAergic populations which have been shown to regulate cortical activation in wakefulness, attention, and learning (Anaclet et al., 2015; Lin, Brown, Hussain Shuler, Petersen, & Kepecs, 2015; Xu et al., 2015; Yang, Thankachan, McCarley, & Brown, 2017).

#### Caudate putamen nucleus

In mice, the combined CPu comprises much of the dorsal striatum and is primarily responsible for integrating cortical and sensory information and enabling regulation of voluntary movement (Schroder, Moser, & Huggenberger, 2020). V5^+^ neurons in the CPu are organized in a lattice-like pattern, where cell bodies are generally distributed at similar distances from each other in three dimensions. tdTomato^+^ branching processes were observed to fill the spaces between these *mWake*^+^ cell bodies (Figure 7g-i). This pattern is seen throughout the CPu, bound by the nucleus accumbens on the medial side, with a somewhat higher density along the cc at the dorsolateral edge, the region more strongly associated with sensorimotor integration rather than reward (Voorn, Vanderschuren, Groenewegen, Robbins, & Pennartz, 2004).

#### Subparafascicular nucleus

Towards the caudal edge of the thalamus, near the junction with the ZI, *mWake^+^* neurons label the parvocellular section of the SPFp (Figure 7j-l). The SPFp is implicated in receiving stimuli from genito-sensory inputs, as well as visual and auditory sensory inputs (Coolen, Veening, Wells, & Shipley, 2003). *mWake* mRNA expression and V5 protein signal was observed in cell bodies within the nucleus. tdTomato^+^ projections were enriched in the lateral aspect of the nucleus and also comprise columnar tracts underlying the optic tract (ot).

#### Subfornical organ

A particularly interesting structure containing *mWake*-expressing cells is the SFO (Figure 7m-o). This circumventricular organ is located along the dorsal edge of the 3v and lacks an effective blood brain barrier due to the presence of fenestrated capillaries. This structural feature enables both osmoregulation and monitoring of hunger and thirst hormones by the SFO (McKinley et al., 2019). mWAKE-V5 signal and *mWake* mRNA label cell bodies in the SFO, and can also be observed in the ependymal cells lining the 3v (see below, Figure 13) (Del Bigio, 2010).

#### Limbic system

We observed *mWake* expression in multiple interconnected structures of the limbic system—the amygdala, lateral septal center (LSc), bed nucleus of the stria terminals (BNST), and islands of Calleja (ICj). These integrate external and internal information to regulate emotion and motivation, as well as downstream effects on behavior and memory (Catani, Dell’acqua, & Thiebaut de Schotten, 2013; Sokolowski & Corbin, 2012). The amygdala produces emotional salience, which is an important driving force behind many behaviors in humans and rodents (Sokolowski & Corbin, 2012). We observed *mWake* mRNA, staining for the mWAKE-V5 fusion protein, and tdTomato fluorescence in several subdivisions of the amygdala (Figure 8a-c). There is a prominent cluster of both *mWake*^+^ cell bodies and processes in the lateral amygdala (LA), with the highest density in the dorsal-most boundary where the external capsule (ec) and cc meet, a region implicated in the acquisition of associative fear learning and its memory storage (Johnson et al., 2008). In the central amygdala (CeA), cell bodies containing *mWake* mRNA and V5 protein and processes expressing tdTomato line its lateral portion along the ec. An additional cluster of cell bodies containing ISH and V5 signal, along with tdTomato fluorescent projections, is found in the posterior portion of the basomedial amygdala nucleus (BMA) (Figure 8a-c). Both the CeA and BMA have been implicated in acquisition of conditioned defensive responses (Fadok, Markovic, Tovote, & Lüthi, 2018).

**Figure 8.**
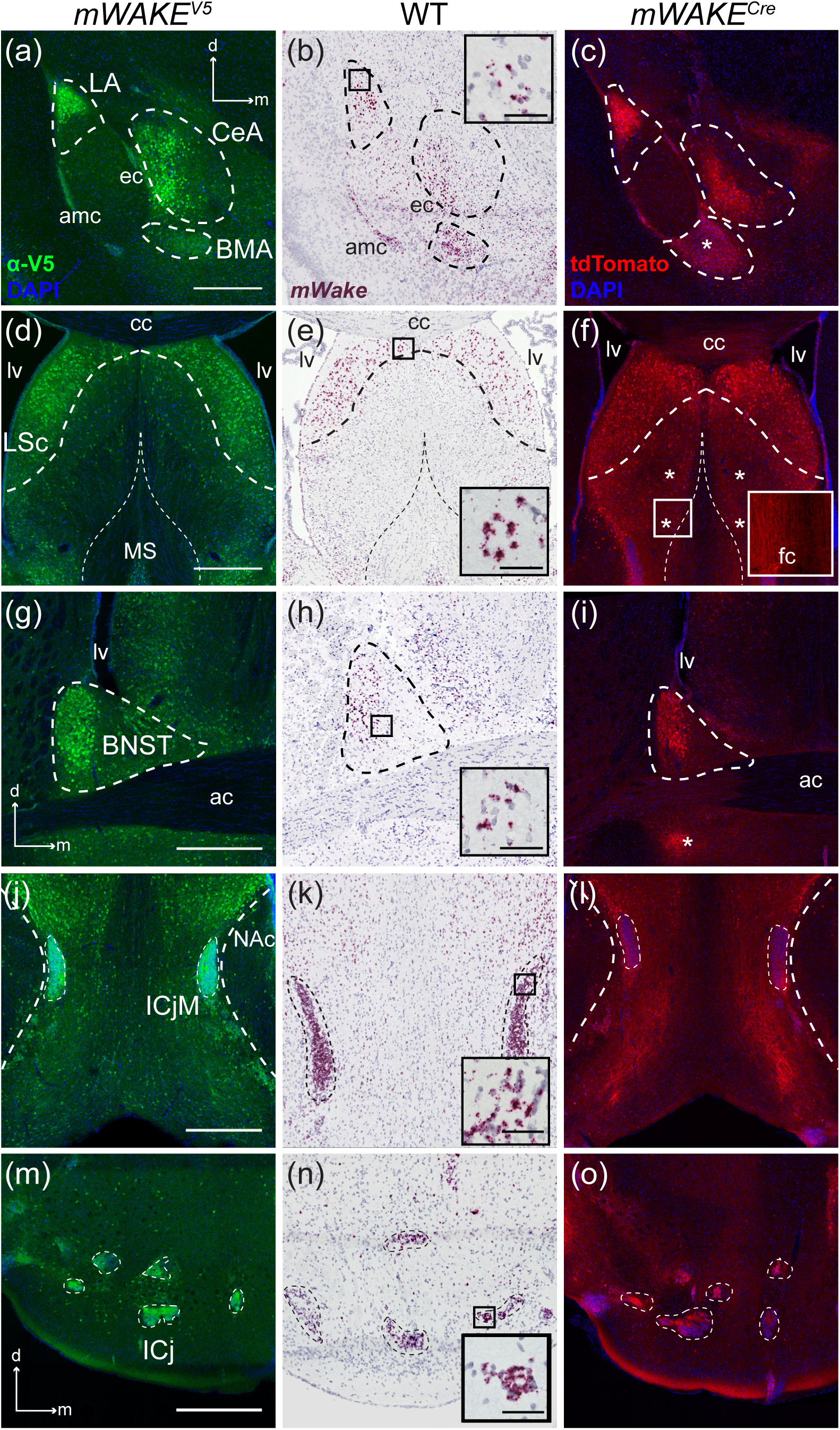
Limbic system centers with *mWake^+^* cells. Representative images of *mWake* expression in brain regions of the limbic system. (a-c) Images of an amygdala area, with sub-regions LA, CeA, and BMA denoted by dashed outlines. (d-f) Images of the LSc, with the thicker dashed outline marking the boundary of the LSc, and the thinner dashed line marking the limits of the MS. (g-i) Images of a BNST, encircled by a dashed line. (j-l) Images of the ICjM, encircled by the thinner dashed line, while the thicker dashed line marks the medial limits of the NAc. (m-o) Images of multiple ICjs, with each individual island outlined by thin dashed lines. (a), (d), (g), (j), and (m) Anti-V5 IF staining (green) with DAPI in *mWAKE^(V5/+)^* mice. (b), (e), (h), (k), and (n) Chromogenic RNAscope ISH for *mWake* mRNA (red) with hematoxylin counterstain (purple) in wild-type (WT) mice. (c), (f), (i), (l), and (o) endogenous tdTomato fluorescence (red) in *mWAKE^(Cre/+)^* mice with DAPI. The black boxes in (b), (e), (h), (k), and (n) depict the approximate location of the representative higher-magnification inset, where the scale bar denotes 50 μm. The white box in (f) indicates the approximate location of the representative higher-magnification inset in the lower right corner. Asterisks in (f) and (i) mark the presence of tdTomato^+^ fibers and processes. Scale bars in (a), (d), (g), (j), and (m) denote 500 μm and apply to other main panels across a row. Arrows in (a), (g), and (m) indicate the dorsal (d) and medial (m) directions, and apply to (b) and (c), (h) and (i), and (n) and (o), respectively. Abbreviations: ac, anterior commissure; amc, amygdalar capsule; BMA, basomedial amygdala; BNST, bed nucleus of the stria terminalis; cc, corpus collosum; CeA, central amygdala; ec, external capsule; ICj, islands of Calleja; ICjM, islands of Calleja, insula magna; LA, lateral amygdala; lv, lateral ventricle; MS, medial septal nucleus; NAc, nucleus accumbens.

The LSc is broadly involved in regulation of both mood and motivation, acting as a primary relay station within the mesocorticolimbic dopaminergic system as well as other limbic areas (Sheehan, Chambers, & Russell, 2004). Here, we observed a dense band of cell bodies containing *mWake* ISH and V5 labeling, under the cc and stretching ventrally adjacent to the lateral ventricles, with tdTomato signal filling the dorsolateral portion of the septal area (Figure 8d-f). This group of cell bodies terminates above the nucleus accumbens (NAc) and ICj. There are additional cell bodies sparsely distributed throughout the LSc, but few in the medial region, where dense tdTomato^+^ projections pass broadly through the septal area and outline the lateral structure of the nDB below. The dorsolateral edge of the LSc, where mWAKE appears to be expressed, has been shown to receive significant catecholaminergic innervation (Lindvall & Stenevi, 1978).

mWAKE is also expressed in the BNST, another limbic region which serves as a major output of the amygdala to integrate mood information with ascending internal-state information from the hypothalamus (Crestani et al., 2013; Lebow & Chen, 2016). In this region, ISH and V5 staining revealed clusters of *mWake^+^* cell bodies in the anterolateral BNST, which is ventrolateral to the lateral ventricles (lv) and considered to modulate anxiety behaviors and fear conditioning (Duvarci, Bauer, & Paré, 2009) (Figure 8g-i). Additionally, a small but dense cluster of tdTomato^+^ projections sit directly underneath the anterior commissure (ac) in the anteroventral portion of the BNST, which receives extensive medullary noradrenergic innervation (Figure 8i) (Gungor & Paré, 2016).

The ICj are small clusters of granule cells and interneurons dispersed throughout the olfactory tubercle. These clusters form bidirectional connections with both ascending dopaminergic circuits and olfactory components of the limbic system (De Marchis et al., 2004; Fallon, Riley, Sipe, & Moore, 1978; Hsieh & Puche, 2013). The density and small size of the ICj cells results in strong and distinct signals from the ISH of *mWake* mRNA, as well as mWAKE-V5 immunolabeling and tdTomato fluorescence, with each of the smaller islands and the larger insula magna (ICjM) containing >90% *mWake^+^* neurons (Figure 8j-o). The presence of *mWake*^+^ cells in multiple components of the limbic system suggests that mWAKE may serve a functional role related to emotional arousal.

### mWAKE expression in the brainstem

#### Midbrain

*mWake* expression was noted in a number of regions within the brainstem, specifically within the midbrain and pons. In the midbrain, we observed a cluster of neurons with signals for *mWake* mRNA, as well as V5 and tdTomato proteins, throughout the ventrolateral periaqueductal grey (vlPAG), with the medial-most cells overlapping the dorsolateral extensions of the dorsal raphe (DR) (Figure 9a-c). ISH-and V5-positive cell bodies are primarily limited to the rostral half of the vlPAG, with few or no cell bodies ventral to the aqueduct as it enlarges further along the caudal axis.

**Figure 9.**
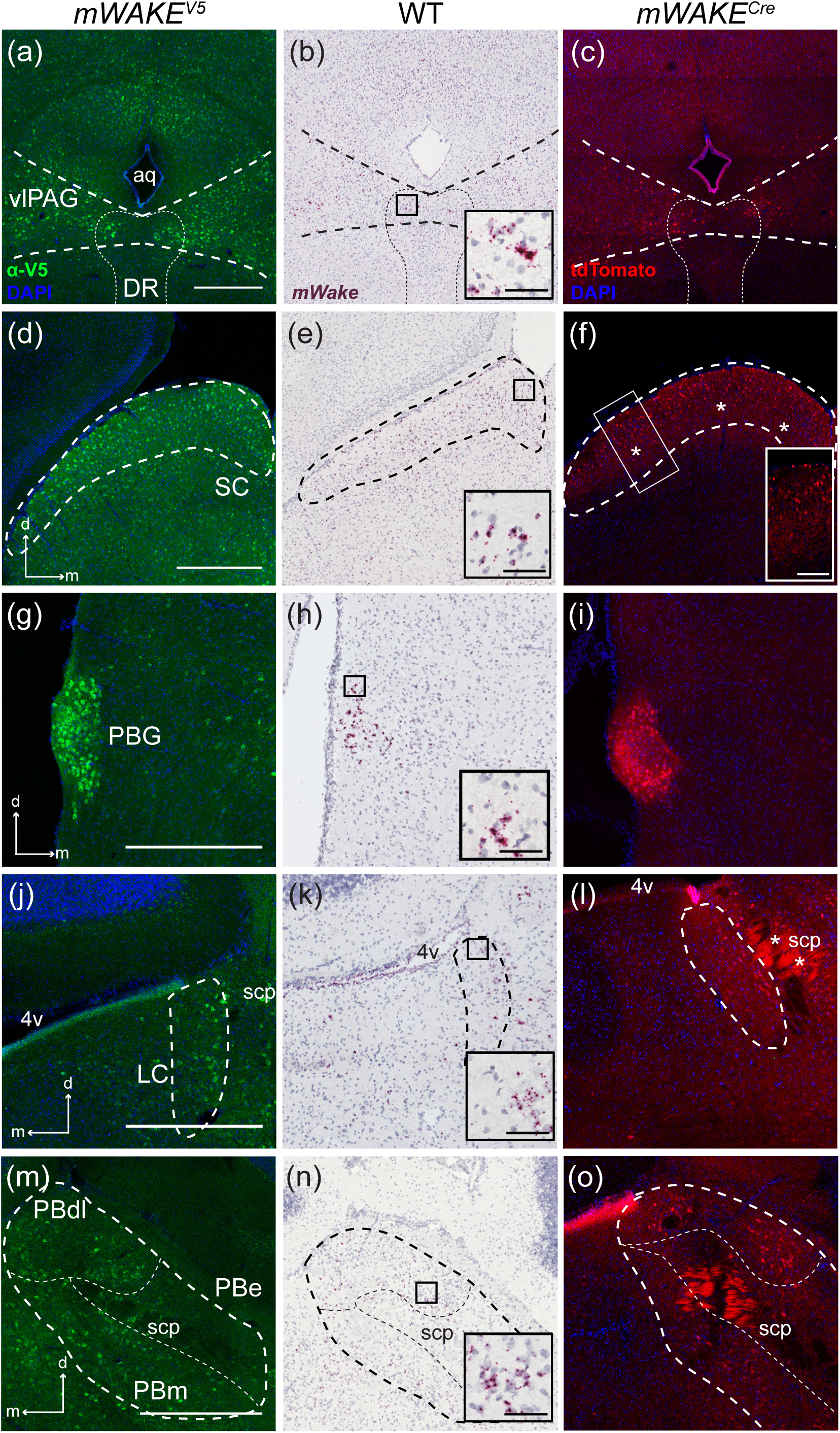
*mWake* expression in brainstem areas. Representative images of *mWake* expression in the brainstem. (a-c) vlPAG denoted by thicker dashed lines, and DR outlined by a thinner dashed line. (d-f) SC, outlined by the dashed line. (g-i) PBG. (j-l) LC area encircled by a dashed line. (m-o) PB, with the entire area encircled by the thicker dashed line, and the PBdl, PBe, and PBm subregions outlined by thin dashed lines. (a), (d), (g), (j), and (m) Anti-V5 IF staining (green) with DAPI in *mWAKE^(V5/+)^* mice. (b), (e), (h), (k), and (n) Chromogenic RNAscope ISH for *mWake* mRNA (red) with hematoxylin counterstain (purple) in wild-type (WT) mice. (c), (f), (i), (l), and (o) Endogenous tdTomato fluorescence (red) in *mWAKE^(Cre/+)^* mice with DAPI. The black boxes in (b), (e), (h), (k), and (n) depict the approximate location of the representative higher-magnification inset in the lower right corner, where the scale bar denotes 50 μm. The white box in (f) indicates the approximate location of the representative higher-magnification inset in the lower right corner. Asterisks in (f) and (l) label tdTomato^+^ fibers. Scale bars in (a), (d), (g), (j), and (m) all denote 500 μm and apply to other main panels across a row. Arrows in (d), (g), (j), and (m) indicate the dorsal (d) and medial (m) directions, and apply to (e) and (f), (h) and (i), (k) and (l), and (n) and (o), respectively. Abbreviations: 4v, fourth ventricle; aq, aqueduct; DR, dorsal raphe; LC, locus coeruleus; PBG, parabigeminal nucleus; PBdl, dorsolateral parabrachial nucleus; PBe, external parabrachial nucleus; PBm, medial parabrachial nucleus; scp, superior cerebellar peduncle; vlPAG, ventrolateral periaqueductal gray.

Along the tectum of the mammalian midbrain, the dorsal-most portion of the superior colliculus (SC) contains a band of cell bodies, as visualized by *mWake* mRNA ISH and V5 staining, as well as significant tdTomato^+^ fibers (Figure 9d-f). This superficial layer of the SC receives afferent projections from both retinal ganglion cells (RGCs) and the visual cortex, which meet in the visually-mapped laminae of the SC, enabling tuning of responses to sensorimotor stimuli (Ito & Feldheim, 2018; Zhao, Liu, & Cang, 2014). Interestingly, the SC is one of the few regions with significant connectivity differences between mice and primates, with only 10% of RGCs projecting to the SC in primates compared to 80% in mice, although there is no evidence that the SC is composed of different cellular and neuronal populations between species (Dhande & Huberman, 2014; Ito & Feldheim, 2018). In addition, the parabigeminal nucleus (PBG) is generally considered a satellite region of the superficial SC. The PBG exhibits dense labeling of *mWake^+^* neurons as demonstrated by *mWake* ISH and V5 staining, as well as interwoven tdTomato^+^ processes (Figure 9g-i) (Graybiel, 1978).

#### Pons

In the pons, *mWake* mRNA- and mWAKE-V5-expressing cells cluster beneath the lateral edge of the 4^th^ ventricle (4v), with groups on both the medial and lateral side of the fiber tract of the superior cerebellar peduncle (scp), and substantial tdTomato^+^ projections stretching across the scp. The most medial of these cells intermingle with the lateral portion of the noradrenergic LC region, which regulates arousal, attention, and wakefulness (Figure 9j-l) (Bell et al., 2020; Schwarz & Luo, 2015). Other mWAKE-V5*^+^* cell bodies and tdTomato^+^ projections located around the scp and anterior to the LC are located within the parabrachial nucleus (PB) (Figure 9m-o). Specifically, *mWake* expression labels both the dorsolateral and medial PB regions, which can produce a persistent coma-like state in rats when lesioned (Fuller, Sherman, Pedersen, Saper, & Lu, 2011). In contrast, there is a distinct absence of *mWake* mRNA, mWAKE-V5, and tdTomato protein signal in the external lateral subsection of the PB, involved in the hypercapnic response to noxious stimuli (Carter, Soden, Zweifel, & Palmiter, 2013; Kaur et al., 2013).

We observed additional *mWake* signals in multiple regions of the brainstem involved in sensory processing and integration. In the auditory-processing cochlear nucleus (CN), *mWake^+^* cells, as visualized by V5 immunostaining and RNAscope, appear to occupy both the ventral and dorsal subdivisions, which are involved in identifying sound localization and source, but it is unclear if *mWake^+^* cells are limited to any particular layer within the CN (Figure 10a-c) (Mao, Montgomery, Kubke, & Thorne, 2015; May, 2000; Shore & Zhou, 2006). *mWake* mRNA, V5 staining, and tdTomato fluorescence also indicate the presence of *mWake^+^* neurons and their projections in the inferior colliculus (IC), the main midbrain structure for processing aural information communicated between the sensory receptive regions and the auditory cortex (Figure 10d-f) (Aitkin, 1986). These cell bodies are primarily limited to the medial aspect of the IC, with tdTomato^+^ projections enriched along the tectum.

**Figure 10.**
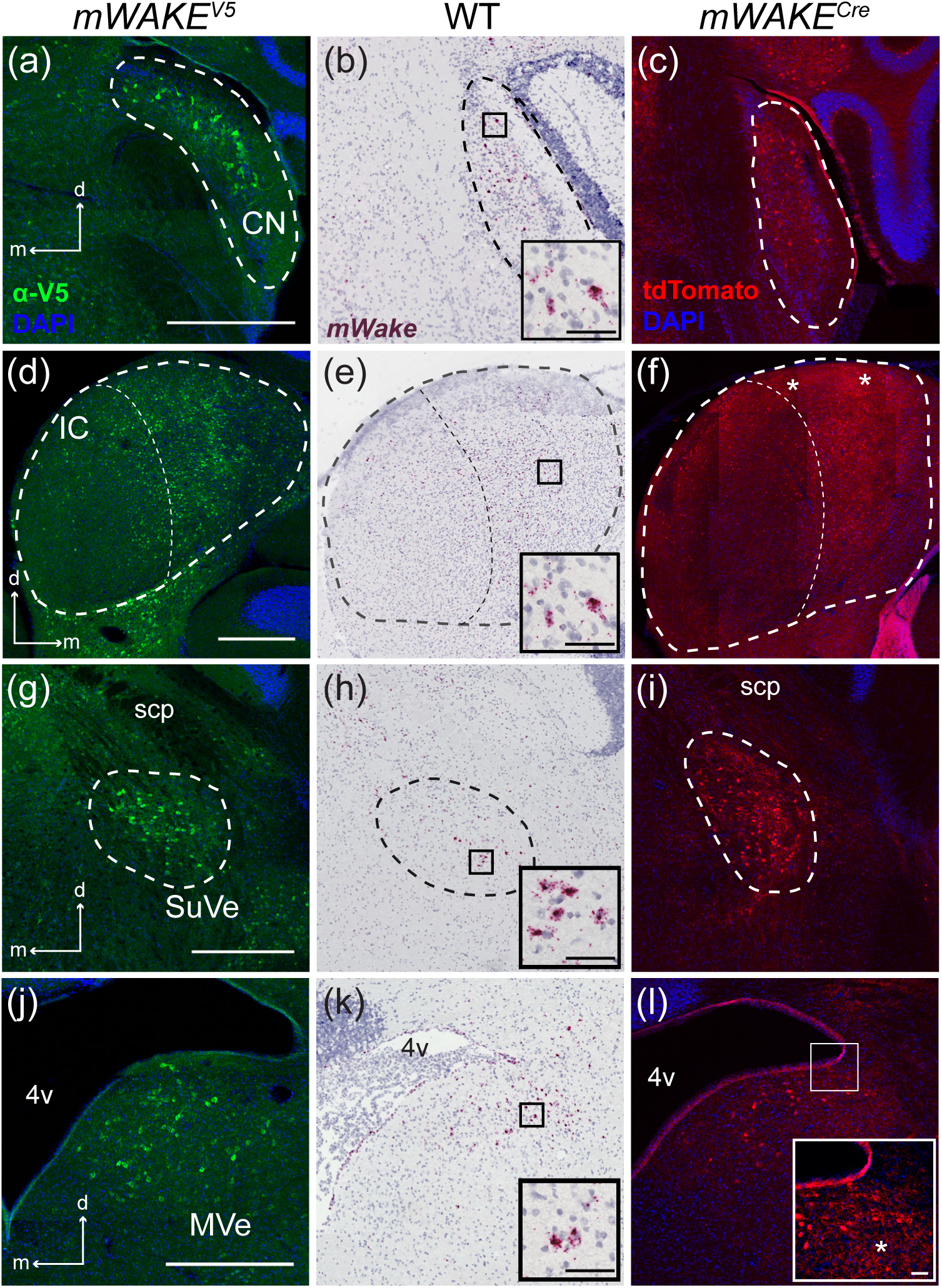
Sensory-related regions containing *mWake* expression. Representative images of *mWake* expression in sensory-related areas of the brainstem. Images of a CN (a-c), IC (d-f), SuVe (g-i), and MVe (j-l) are shown. In (a-c), the dashed line outlines the CN. In (d-f), the thicker dashed line outlines the IC, which is divided into medial and lateral portions by the thinner dashed line. In (g-i), the dashed line encircles the SuVe. (a), (d), (g), and (j) Anti-V5 IF staining (green) with DAPI in *mWake^(V5/+)^* mice. (b), (e), (h), and (k) Chromogenic RNAscope ISH for *mWake* mRNA (red) with hematoxylin counterstain (purple) in wild-type (WT) mice. (c), (f), (i), and (l) endogenous tdTomato fluorescence (red) in *mWake^(Cre/+)^* mice with DAPI. The black boxes in (b), (e), (h), and (k) depict the approximate location of the representative higher-magnification insets in the lower right corner, where the scale bar denotes 50 μm. The white box in (l) indicates the approximate location of the representative higher-magnification inset in the lower right corner. Asterisks in (f) and (l) mark tdTomato^+^ fibers. Scale bars in (a), (d), (g), and (j) denote 500 μm and apply to other main panels across a row. Arrows in (a), (d), (g) and (j) indicate the dorsal (d) and medial (m) directions, and apply to other main panels across a row. Abbreviations: 4v, fourth ventricle; CN, cochlear nucleus; IC, inferior colliculus; MVe, medial vestibular nucleus; scp, superior cerebellar peduncle; SuVe, superior vestibular nucleus.

We further observed two anatomically distinct populations of *mWake*-expressing neurons within the brainstem vestibular nuclei, involved in kinesthesia and the maintenance of balance. *mWake* mRNA and mWAKE-V5 protein marked cell bodies found clustered in the superior vestibular nucleus (SuVe) directly under the scp (Figure 10g-i). Similarly, signals from *mWake*^+^ cell bodies are scattered in the larger medial vestibular nucleus (MVe), directly caudal to the SuVe, where we observed tdTomato^+^ processes innervating much of the area directly ventral to the lateral wing of the 4v (Figure 10j-l).

### Cortical regions with *mWake*^+^ cells

In addition to subcortical and brainstem expression, mWAKE is observed in several structures in both the paleocortex and neocortex. In the olfactory transduction pathway, the olfactory bulb (OB) serves as the primary relay, receiving inputs directly from receptor neurons of the olfactory endothelium and transmitting scent information to the olfactory cortex and tubercle for processing (Nagayama, Homma, & Imamura, 2014). We observed substantial labeling of *mWake^+^* cell bodies, as evidenced by ISH and V5 staining, in the granule layer of the OB, which is comprised mostly of inhibitory interneurons (Figure 11a,b). However, there are also extensive tdTomato^+^ processes innervating the superficial glomeruli layer, where mitral and tufted cells synapse with the primary olfactory receptor axons, suggesting a role for *mWake^+^* neurons in the initial processing of odor sensation (Figure 11c) (Nagayama, Homma, & Imamura, 2014).

**Figure 11.**
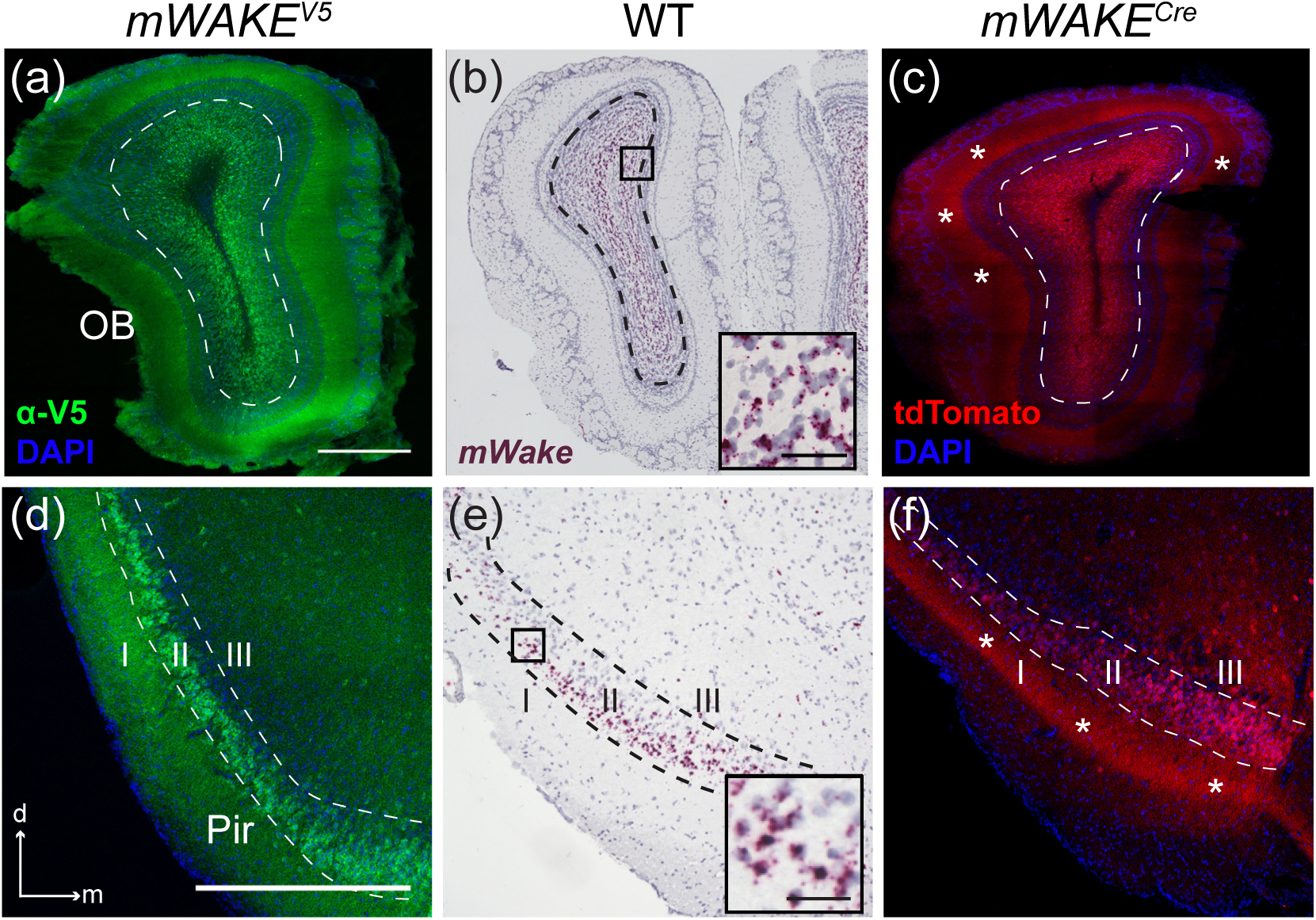
*mWake^+^* expression in the paleocortex. Representative images of *mWake* expression in 2 structures of the paleocortex. (a-c) OB, with dashed line indicating the outer limits of the granule cell layer. (d-f) Pir, with dashed lines separating layers I, II, and II. (a) and (d) Anti-V5 IF staining (green) with DAPI in *mWAKE^(V5/+)^* mice. (b) and (e) Chromogenic RNAscope ISH for *mWake* mRNA (red) with hematoxylin counterstain (purple) in wild-type (WT) mice. (c) and (f) Endogenous tdTomato fluorescence (red) in *mWAKE^(Cre/+)^* mice with DAPI. In (c) and (f), asterisks indicate tdTomato^+^ processes. Black boxes in (b) and (e) depict the approximate location of the representative higher-magnification inset in the lower right corner, where the scale bar denotes 50 μm. Scale bars in (a) and (d) denote 500 μm, and apply to other main panels across a row. Arrows in (d) indicate the dorsal (d) and medial (m) directions, and apply to (e) and (f). Abbreviations: OB, olfactory bulb; Pir, piriform cortex.

The piriform area (Pir) contains one of the densest layers of *mWake^+^* cell bodies seen throughout the brain. As shown by anti-V5 labeling, ISH, and tdTomato fluorescence, these neurons broadly occupy pyramidal layer II, with tdTomato^+^ projections extending outwards through the superficial layer I, as well as inwards through the less dense layer III (Figure 11d-f). Pyramidal cells of Pir layer II have been shown to synapse with granule cells of the OB and ICj, as well as with multiple olfactory subcircuits of the amygdala, to process and integrate olfactory information with memory and emotional salience (Bekkers & Suzuki, 2013; Illig & Wilson, 2009; Soudry, Lemogne, Malinvaud, Consoli, & Bonfils, 2011).

In the neocortex, *mWake* expression is restricted to several discrete locations. Throughout much of the somatosensory (SS) cortex, including the primary (SSp), barrel field (SSb), and supplemental areas (SSs), cell bodies containing *mWake* mRNA and V5 protein label layer IV, while tdTomato-filled processes extend through the more superficial layers I, II, and III (Figure 12a-d). This thin band of cell bodies is nearly absent in layer IV of the more caudal parietal and auditory areas. Although the morphology of *mWake^+^* cells is not easily discerned in these dense bands, most of layer IV of the SSp are comprised of stellate cells (∼75%), rather than the pyramidal neurons which comprise the majority in other cortical areas (Scala et al., 2019). The auditory (AudC) and temporal (TeA) cortical areas contain dense labeling of neurons with *mWake* mRNA, V5 protein, and tdTomato which nearly fill layers II/III (Figure 12e-g). The anterior-posterior distribution of this cortical cluster is delimited anteriorly by the visceral and gustatory cortical areas, and posteriorly by the entorhinal cortical areas. Within the central sulcus above the medial transmission of the cc, the anterior cingulate cortex (ACC) has been implicated as a central node for cognitive processes involving motivation and as a hub for social interactions (Apps, Rushworth, & Chang, 2016; Holroyd & Yeung, 2012). *mWake*^+^ cell bodies are observed in layer IV of the ventral portion of this structure, with processes tagged with tdTomato extending medially (toward the central sulcus) through the superficial layers (Figure 12h-j). In addition to the *mWake^+^* cortical cell bodies, the tdTomato fluorophore labels the cc underneath these cortical regions, suggesting that axons emanating from *mWake^+^* neurons are present in transhemispheric and subcortical-cortical axonal pathways (Figure 12k).

**Figure 12.**
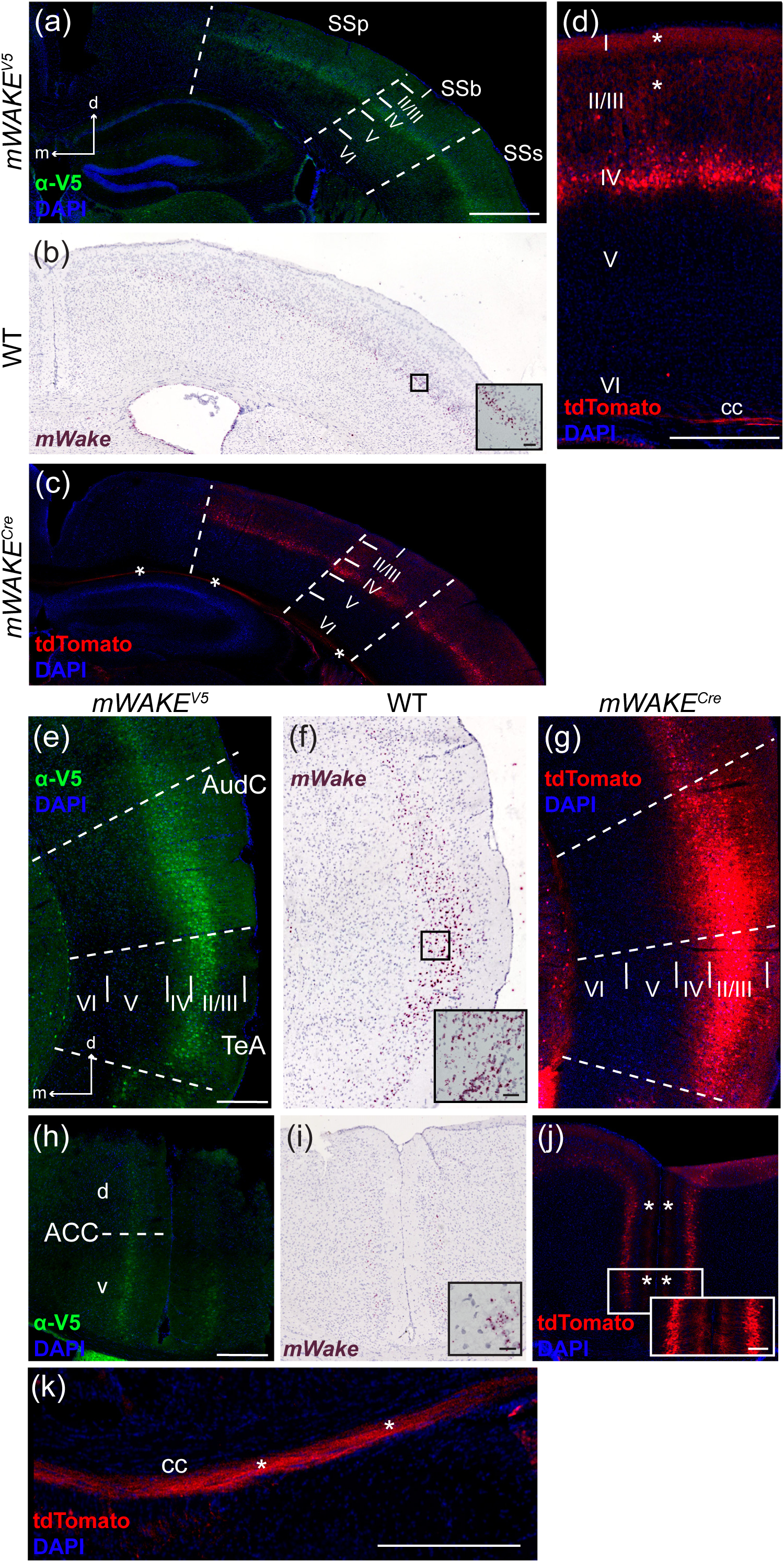
*mWake* expression within the neocortex. Representative images of *mWake* expression in the neocortex. (a-c) Somatosensory area of the cortex, with dashed lines separating the SSp, SSb, and SSs, and solid lines delineating layers I, II/III, IV, V, and VI. (d) Higher-magnification image of the SSb area containing tdTomato^+^ cell bodies and processes, with cortical layers labeled. (e-g) AudC and TeA, separated by dashed lines with cortical layers I, II/III, IV, V, and VI separated by solid lines. (h-j) Bilateral ACC, with a dashed line separating the dorsal (d) and ventral (v) portions. (k) Higher-magnification image of tdTomato fluorescence in the cc. (a), (e), and (h) Anti-V5 IF staining (green) with DAPI in *mWAKE^(V5/+)^* mice. (b), (f), and (i) Chromogenic RNAscope ISH for *mWake* mRNA (red) with hematoxylin counterstain (purple) in wild-type (WT) mice. (c), (d), (g), (j), and (k) Endogenous tdTomato fluorescence (red) in *mWAKE^(Cre/+)^* mice with DAPI. In (c), (d), (j), and (k), asterisks indicate tdTomato^+^ processes. The black boxes in (b), (f), and (i) depict the approximate location of the representative higher-magnification inset in the lower right corner, where the scale bar denotes 50 μm. Scale bar in (a) denotes 1 mm, and applies to (b) and (c). Scale bars in (d) and (k) denote 300 μm. Scale bars in (e) and (h) denote 500 μm and apply to (f-g) and (i-j), respectively. Arrows in (a) and (e) indicate the dorsal (d) and medial (m) directions, and apply to (b), (c), (f) and (g). Abbreviations: ACC, anterior cingulate cortex; AudC, auditory cortex; cc, corpus collosum; SSb, somatosensory cortex barrel region; SSp, primary somatosensory cortex; SSs, supplemental somatosensory cortex; TeA, temporal cortex area.

### Ependymal cells

mWAKE expression is not solely restricted to neurons. mWAKE-V5, *mWake* mRNA, and tdTomato protein were also found in ependymal cells, which form the walls of the ventricular system and regulate access between the brain parenchyma and the cerebrospinal fluid (CSF) (Jimenez, Dominguez-Pinos, Guerra, Fernandez-Llebrez, & Perez-Figares, 2014) (Figure 13a-i). In the hypothalamus, we observed *mWake* mRNA, as well as anti-V5 and tdTomato signal, in these cells lining the dorsal portion of the 3v (Figure 13a-c), and they constitute a significant cluster in our hypothalamic scRNAseq data, identified by high levels of *Foxj1* expression (Figure S3). However, it is worth noting that these ependymal cells are likely overrepresented in our scRNA-Seq dataset, since they tend to survive the sorting procedure better than neurons (Saxena et al., 2012). In addition to the 3v, we noted similar expression of *mWake* mRNA, mWAKE-V5, and tdTomato in ependymal cells lining the lv and 4v (Figure 13d-i). These findings are somewhat reminiscent of the expression of mWAKE in the SFO (Figure 7m-o), which together suggest a potential role for mWAKE in osmoregulation.

**Figure 13.**
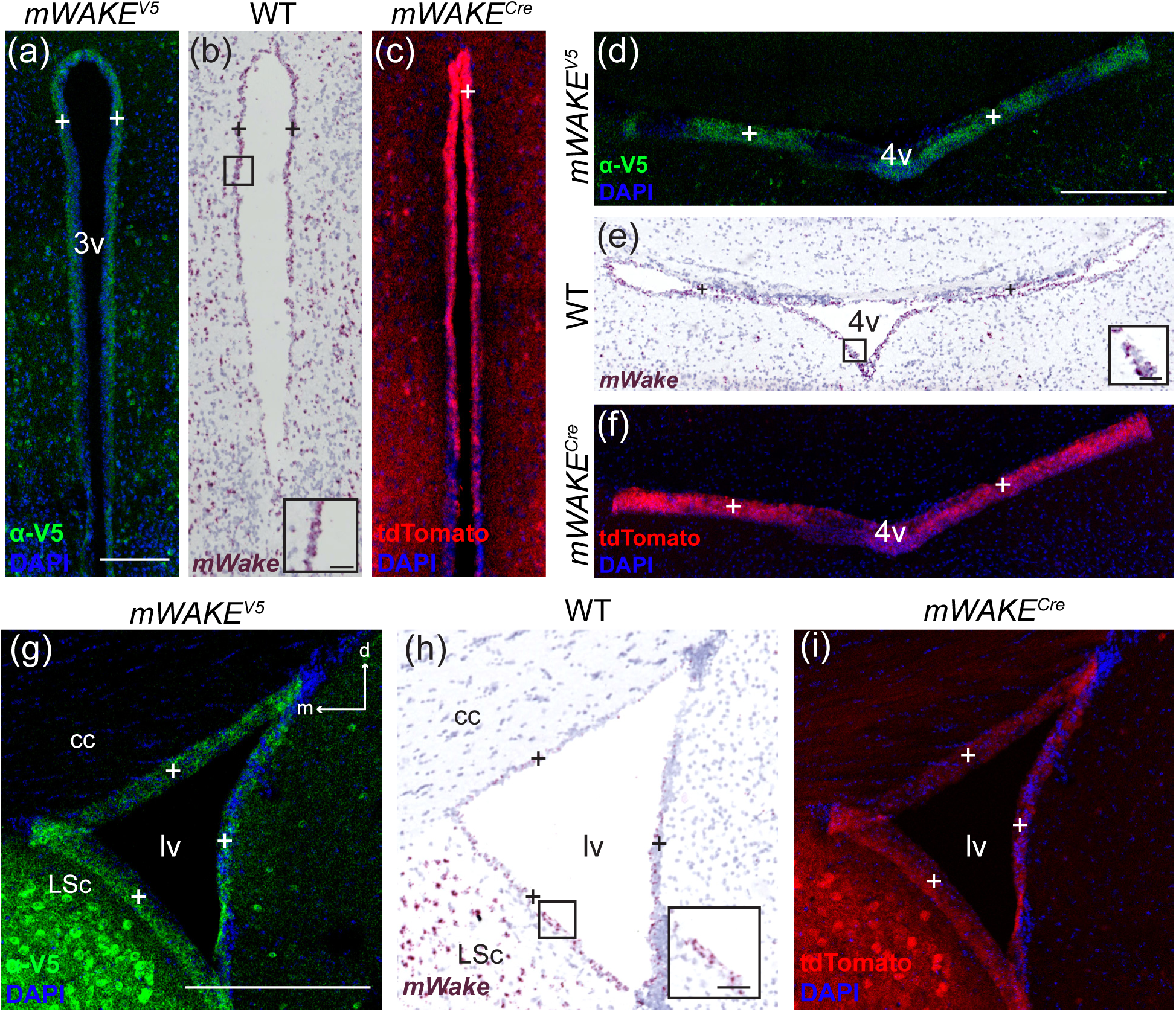
Ependymal cells express *mWake*. Representative images of *mWake* expression in ependymal cells. Images of the 3v (a-c), 4v (d-f), and one lv (g-i) are shown. (a), (d), and (g) Anti-V5 IF staining (green) with DAPI in *mWAKE^(V5/+)^* mice. (b), (e), (h) Chromogenic RNAscope ISH for *mWake* mRNA (red) with hematoxylin counterstain (purple) in wild-type (WT) mice. (c), (f), and (i) Endogenous tdTomato fluorescence (red) in *mWAKE^(Cre/+)^* mice with DAPI. In all images, “+” indicates *mWake^+^* ependymal cells. Black boxes in (b), (e), and (h) depict the approximate location of the representative higher-magnification inset in the lower right corner, where the scale bar denotes 50 μm. Scale bars in (a), (d), and (g) denote 300 μm and apply to (b and c), (e and f), and (h and i), respectively. Arrows in (g) indicate dorsal (d) and medial (m) directions, and apply to (h) and (i). Abbreviations: 3v, third ventricle; 4v, fourth ventricle; cc, corpus collosum; LSc, lateral septal nucleus; lv, lateral ventricle.

### mWAKE is expressed in undefined structures

To identify regions and structures labelled by mWAKE, we relied on standard atlases in the field: Paxinos and Franklin’s *The Mouse Brain in Stereotaxic Coordinates* and the Allen Brain Institute’s Reference Mouse Brain Atlas. Using these references to the best of our ability, we were able to systematically catalog mWAKE expression, with two prominent exceptions. The first sits at the junction of the hypothalamic and thalamic regions, directly above the 3v and delimited below by the paraventricular nucleus, which lacks mWAKE expression (Figure 14a-c). The thalamic structure in this area, the reuniens nucleus (Re), appears to delineate the region lacking mWAKE seen in our RNAscope, anti-V5, and tdTomato fluorescence imaging. The undefined mWAKE labeling may belong to a caudal extension of the BNST, but we have not found any previously published description of such a structure. The other undefined cluster of *mWake^+^* cells appears at the junction of the paleocortex and neocortex, and forms a round structure, which is tubular in the rostral-caudal axis. This structure is located lateral to the LA (Figure 14d-f). It is possible that this cell group is part of the dorsal endopiriform area, or possibly layer V/VI of the rostral end of the entorhinal cortex; however, the round shape and scattered cell bodies do not readily match the laminar organization of these structures. Although there are relatively few *mWake^+^* cells in this cluster, its location is remarkably consistent between hemispheres and between animals, and it is possible the *mWake^+^* cell bodies label an undefined cortical subregion.

**Figure 14.**
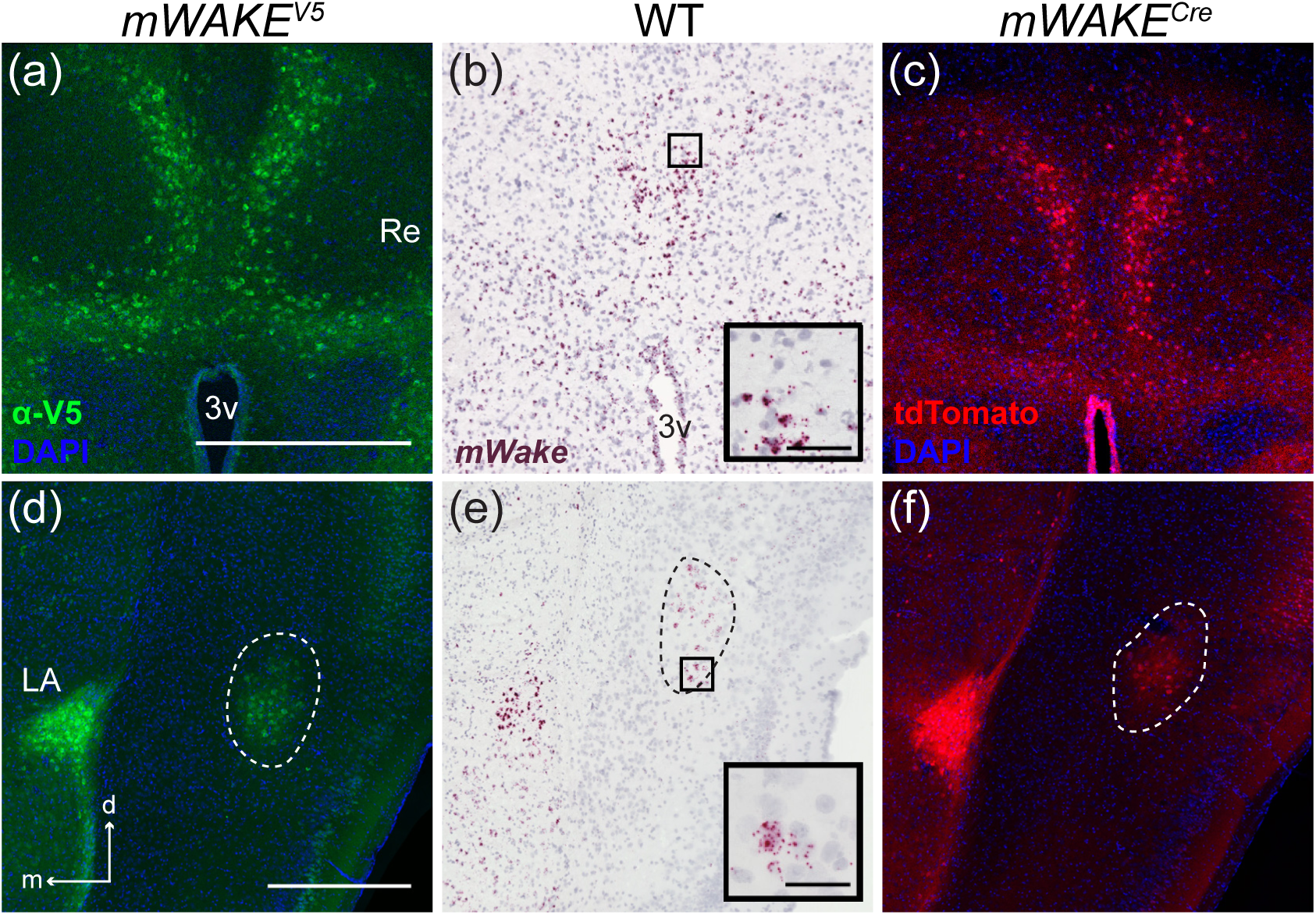
*mWake* expression in undefined regions. Representative images of *mWake* expression in 2 undefined structures. Images of mWAKE expression in supra-hypothalamic protrusions (a-c) and a round mid-cortical cluster (d-f), encircled by a dashed line. (a) and (d) Anti-V5 IF staining (green) with DAPI in *mWAKE^(V5/+)^* mice. (b) and (e) chromogenic RNAscope ISH for *mWake* mRNA (red) with hematoxylin counterstain (purple) in wild-type (WT) mice. (c) and (f) Endogenous tdTomato fluorescence (red) in *mWAKE^(Cre/+)^* mice with DAPI. The black boxes in (b) and (e) depict the approximate location of the representative higher-magnification inset in the lower right corner, where the scale bar denotes 50 μm. The scale bars in (a) and (d) denote 500 μm, and apply to other main panels across a row. Arrows in (d) indicate the dorsal (d) and medial (m) directions, and apply to (e) and (f). Abbreviations: 3v, third ventricle; LA, lateral amygdala; Re, thalamic reuniens nuclei.

## Discussion

In this study, we performed a brain-wide analysis of *mWake* expression using three complementary approaches that allow visualization of *mWake* mRNA, protein, and cellular processes. Overall, we show that *mWake^+^* cell bodies are found in a restricted pattern, but distributed across multiple levels of the neuraxis, including the brainstem, subcortical, and cortical regions. Although *mWake* expression can be observed in specific areas in multiple brain regions, there are locations distinctly lacking mWAKE expression. For example, in the brainstem, there is little to no clear mWAKE expression in the medulla; while we did observe sparse V5-expressing cells within the dorsomedial aspect, it was unclear whether this represented *bona fide* mWAKE expression, as the ISH for *mWake* mRNA and tdTomato reporter did not label this region. In general, the thalamus is largely devoid of *mWake*^+^ neurons, with the exception of SPFp. However, the thalamus is intensely labeled with tdTomato^+^ processes, suggesting signaling influenced by *mWake* passes through and connects to thalamic circuits. Interestingly, despite its presence in several areas with direct connections to the hippocampus (e.g., amygdala, BF), we observed no evidence of *mWake* expression in any components of the hippocampus. *mWake* is also not found in most broad neocortical regions, such as the frontal or occipital lobes, as well as the motor cortex. Moreover, in those cortical regions containing *mWake^+^* cells, they are limited almost exclusively to Layer II/III or IV. In contrast to the discrete labeling of *mWake^+^* cells in specific regions, tdTomato labeling observed in *mWAKE^Cre^* mice was often diffusely distributed throughout much of the brain, suggesting that mWAKE-expressing cells receive or transmit information broadly; this conclusion is supported by our previous projection tracing of *mWake^DMH^* neurons (Bell et al., 2020).

To complement these imaging experiments, we also performed detailed analyses of scRNA-Seq data from hypothalamic *mWake^+^* cells. We chose the hypothalamus because this region contains the highest number of *mWake*-expressing sub-regions throughout the brain and likely contains the highest overall number of *mWake^+^* cells. These analyses reveal the diversity of cell types that express mWAKE and set the stage for future, highly refined dissections of *mWake^+^* sub-circuit function. Of particular interest is the characterization of *mWake* expression in the SCN. Here, *mWake* is largely confined to the classical core region, where it substantially overlaps with *Vip* and *Grp* expression. *Grp*^+^ and especially *Vip*^+^ neurons have been shown to receive direct projections from ipRGCs and are thought to mediate light-sensitive circadian behaviors such as photoentrainment (Jones, Simon, Lones, & Herzog, 2018; Mazuski et al., 2018; Welsh et al., 2010; Yan et al., 2007). Strikingly, this parallels the expression and function of WAKE in the large ventrolateral (l-LNv) neurons in *Drosophila* (Liu et al., 2014). The l-LNvs are photorecipient clock neurons in flies that are weakly rhythmic and send major projections to the small ventrolateral (s-LNv) neurons, which function as master circadian pacemaker neurons in constant darkness (Nitabach & Taghert, 2008). Similarly, in mice the photorecipient *Vip*^+^ and *Grp*^+^ SCNc neurons are weakly rhythmic and project to the strongly rhythmic SCNs neurons (Abrahamson & Moore, 2001; Welsh et al., 2010; Yan et al., 2007). These observations suggest that *mWake* in the SCNc may serve a special role in light-dependent circadian behaviors in mammals, the details of which remain to be elucidated. In addition, *mWake* appears to define an additional subset of SCNc neurons that lack *Vip* and *Grp*, but are instead *Rorb^+^* and *Penk^+^*. This group of previously undescribed neurons are predicted to reside more dorsally in the SCNc than the *Grp^+^* cells, and may comprise a novel SCN subcircuit.

Our previous studies of WAKE in fruit flies and *mWake* in mice (Bell et al., 2020; Liu et al., 2014; Tabuchi et al., 2018) suggest that these molecules act downstream of the circadian clock to upregulate specific receptors or channels and inhibit neural excitability at night. Based on these prior findings, we speculate that *mWake* might specifically label neurons involved in rhythmic processes or behaviors. For example, we have previously shown that, outside of the SCN, *mWake^+^* neurons in the DMH also exhibit cyclical firing, and loss of mWAKE in these cells selectively promotes locomotor activity at night (Bell et al., 2020). Although yet unstudied, it is plausible that a similar situation exists for *mWake^+^* neurons across the brain. Interestingly, while *mWake* generally labels neurons, non-neuronal ependymal cells lining the 3v, lv, and 4v are also largely *mWake^+^*. These glial cells form a partial barrier between the CSF and brain parenchyma and assist with osmoregulation, as well as regulation of transport for various molecules between the brain and CSF compartments (Jimenez et al., 2014). Recently, in *Drosophila*, permeability of the blood-brain barrier has been shown to be under circadian control, likely via specific multidrug transporter molecules (Cuddapah, Zhang, & Sehgal, 2019; Zhang, Yue, Arnold, Artiushin, & Sehgal, 2018). Interestingly, ependymal cells in mammals have been shown to exhibit rhythmic clock gene expression (Guilding, Hughes, Brown, Namvar, & Piggins, 2009; Wen et al., 2020). Thus, the expression of *mWake* in ependymal cells and its role as a clock output molecule raise the possibility that the permeability of the brain/CSF interface in mammals is also modulated by circadian influences, possibly through mWAKE.

In the mouse brain, *mWake* is expressed in a limited number of neurons that are distributed across a variety of regions. Many of these regions can be loosely grouped into different functional systems. For instance, the DMH, TMN, LH, and PB have all been implicated in regulating arousal, while the amygdala, LSc, BNST, and ICj can be considered part of the limbic system. In addition, several regions (e.g., OB, CN, IC, SuVe, MVe) are directly connected to a role in sensory perception and processing. These different systems may be functionally distinct and unrelated. However, the prominent hyperarousal phenotype seen in *mWake* mutants makes it tempting to speculate that these *mWake^+^* circuits may all be tied to the perception of arousal. That is, in addition to “classical” arousal regions, we hypothesize that mWAKE functions in the limbic and sensory systems to enable tuning of “emotional” and “sensory” arousal in a rhythmic manner.

In addition, *mWake* expression is often found in multiple nodes along a circuit pathway. For example, *mWake* labels several components of the limbic system, all of which form a highly interconnected network with bidirectional pathways (Catani et al., 2013; Sokolowski & Corbin, 2012). Furthermore, *mWake* is also found in both the OB and its downstream target (Pir), as well as the CN and its direct (IC) and indirect (AudC) downstream targets. A similar situation exists for *Drosophila* WAKE, which acts at multiple levels of the clock network (i.e., l-LNv and DN1p) (Liu et al., 2014; Tabuchi et al., 2018). Although the function of mWAKE expression in multiple components of a given circuit pathway remains unclear, one possibility may be to amplify rhythmic control of circuit activity. In summary, our systematic characterization of *mWake* expression in the mouse brain lays the foundation for future investigations into the role of this protein in a variety of neural circuits and behaviors. We hypothesize that *mWake^+^* cells are particularly sensitive to intrinsic circadian input and that they regulate light-dependent circadian behaviors, arousal, emotional valence, and sensory perception. Future studies using intersectional approaches with the *mWAKE^Cre^* line should help unravel the specific circuit mechanisms underlying rhythmic modulation of these processes.

## List of Abbreviations

I-VI: layers of the cortex
3v: third ventricle
4v: fourth ventricle
ac: anterior commissure
ACC: anterior cingulate cortex
aq: cerebral aqueduct
AudC: auditory cortex
BF: basal forebrain
BMA: basomedial amygdala
BNST: bed nucleus of the stria terminalis
cc: corpus callosum
CeA: central amygdala
CN: cochlear nucleus
CPu: caudate putamen
DMH: dorsomedial hypothalamus
DR: dorsal raphe
ec: external capsule
fc: fornicular column
fx: fornix
IC: inferior colliculus
ICj: islands of Calleja
ICjM: islands of Calleja insula magna
LA: lateral amygdala
LC: locus coeruleus
LH: lateral hypothalamus
LSc: lateral septal center
lv: lateral ventricle
MS: medial septum
MVe: medial vestibular nucleus
NAc: nucleus accumbens
nDB: nucleus of the diagonal band
OB: olfactory bulb
oc: optic chias
ot: optic tract
PBdl: dorsolateral parabrachial nucleus
PBe: external parabrachial area
PBG: parabigeminal nucleus
PBm: medial parabrachial nucleus
Pir: piriform cortex
POA: preoptic area
Re: thalamic reuniens nucleus
RGC: retinal ganglion cell
SC: superior colliculus
scp: superior cerebellar peduncle
SCN: suprachiasmatic nucleus
SCNc: core region of SCN
SCNs: shell region of SCN
SFO: subfornical organ
SPFp: parvocellular subparafascicular nucleus
SSp: primary somatosensory cortex
SSb: barrel field somatosensory cortex
SSs: supplemental somatosensory cortex
SuVe: superior vestibular nucleus
TeA: temporal cortical area
TMN: tuberomammillary nucleus
vlPAG: ventrolateral periaqueductal gray
VMH: ventromedial hypothalamus
vmPOA: ventromedial preoptic area
ZI: zona incerta

## Acknowledgments

We thank members of the Wu Lab for discussion. This work was supported by NIH grants R01DK108230 (S.B.) and R01NS094571 (M.N.W.) and a NINDS Center grant NS050274 for generation of the *mWAKE^V5^* mouse line and access to confocal microscopes.

---

**Figure S1.**
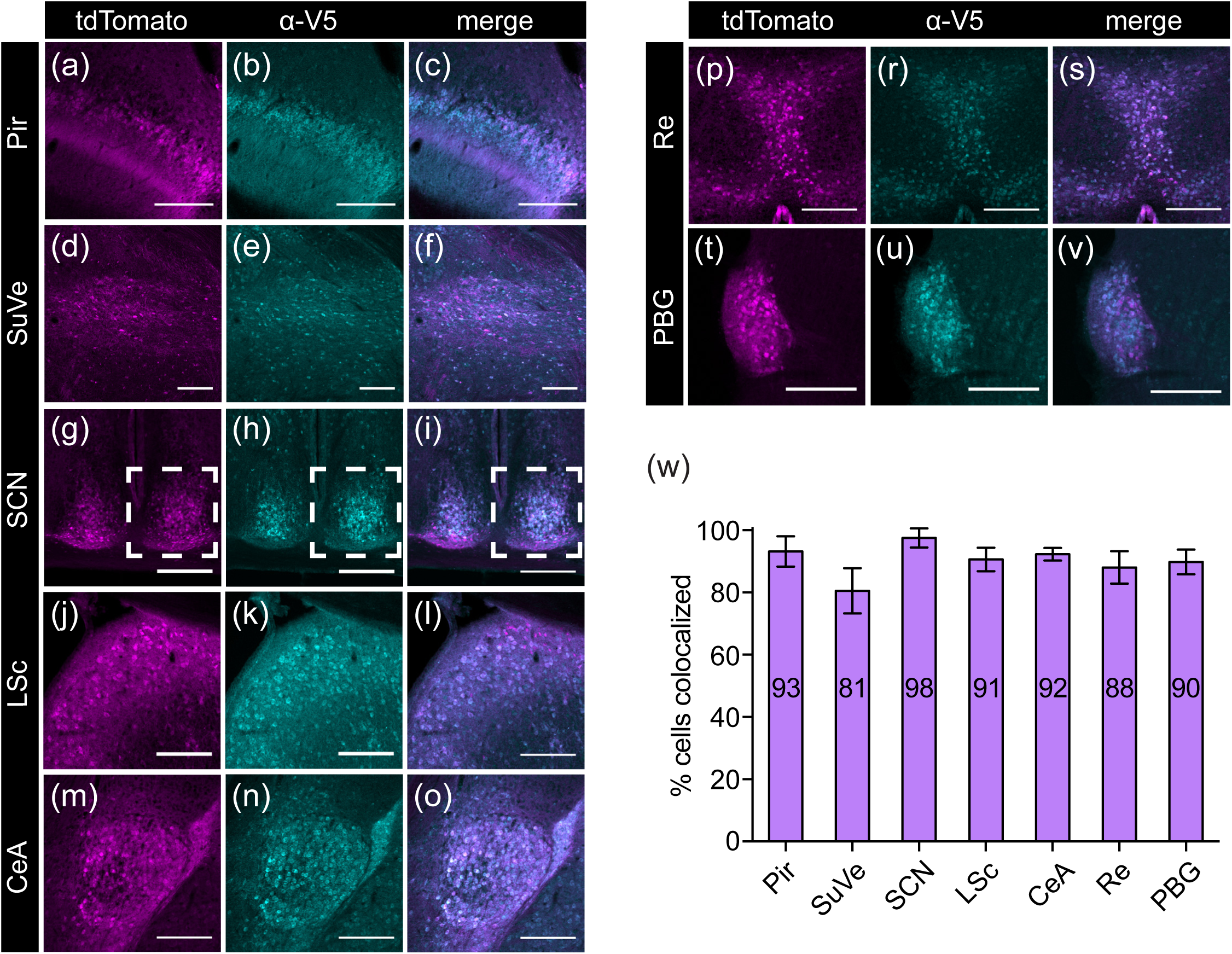
Colabeling of mWAKE-V5 and tdTomato. Images used are representative of 7 locations with discrete *mWake* expression for assessing correlation at the cellular level between *mWAKE^V5^* and *mWAKE^Cre^* transgenic models. Representative images from Pir (a-c), SuVe (d-f), SCN (g-i), LSc (j-l), CeA (m-o), cells adjacent to the Re (p-s), and PBG (t-v) are shown. Dashed outline in (g-i) indicates the unilateral portion used for quantification. (a), (d), (g), (j), (m), (p) and (t) Endogenous tdTomato fluorescence (magenta) in *mWAKE^(V5/Cre)^* mice. (b), (e), (h), (k), (n), (r), and (u) Anti-V5 IF staining (cyan) in the same *mWAKE^(V5/Cre)^* mice. (c), (f), (i), (l), (o), (s), and (v) Merged images for counting colabeled cells (white). (w) Colabeling quantification presented as % of V5^+^ cells with tdTomato fluorescence. Number of tdTomato^+^ cells and V5^+^ cells counted: Pir (n=404; n=423), SuVe (n=251; n=281), SCN (n=428; n=408), LSc (n=429; n=467), CeA (n=431; n=449), near Re (n = 430; n=468), PBG (n=246; n=261). All scale bars represent 200 μm. Abbreviations: CeA, central amygdala; LSc, lateral septal center; PBG, parabigeminal nucleus; Pir, piriform cortex; Re, thalamic reuniens nucleus; SCN, suprachiasmatic nucleus; SuVe, superior vestibular nucleus.

**Figure S2.**
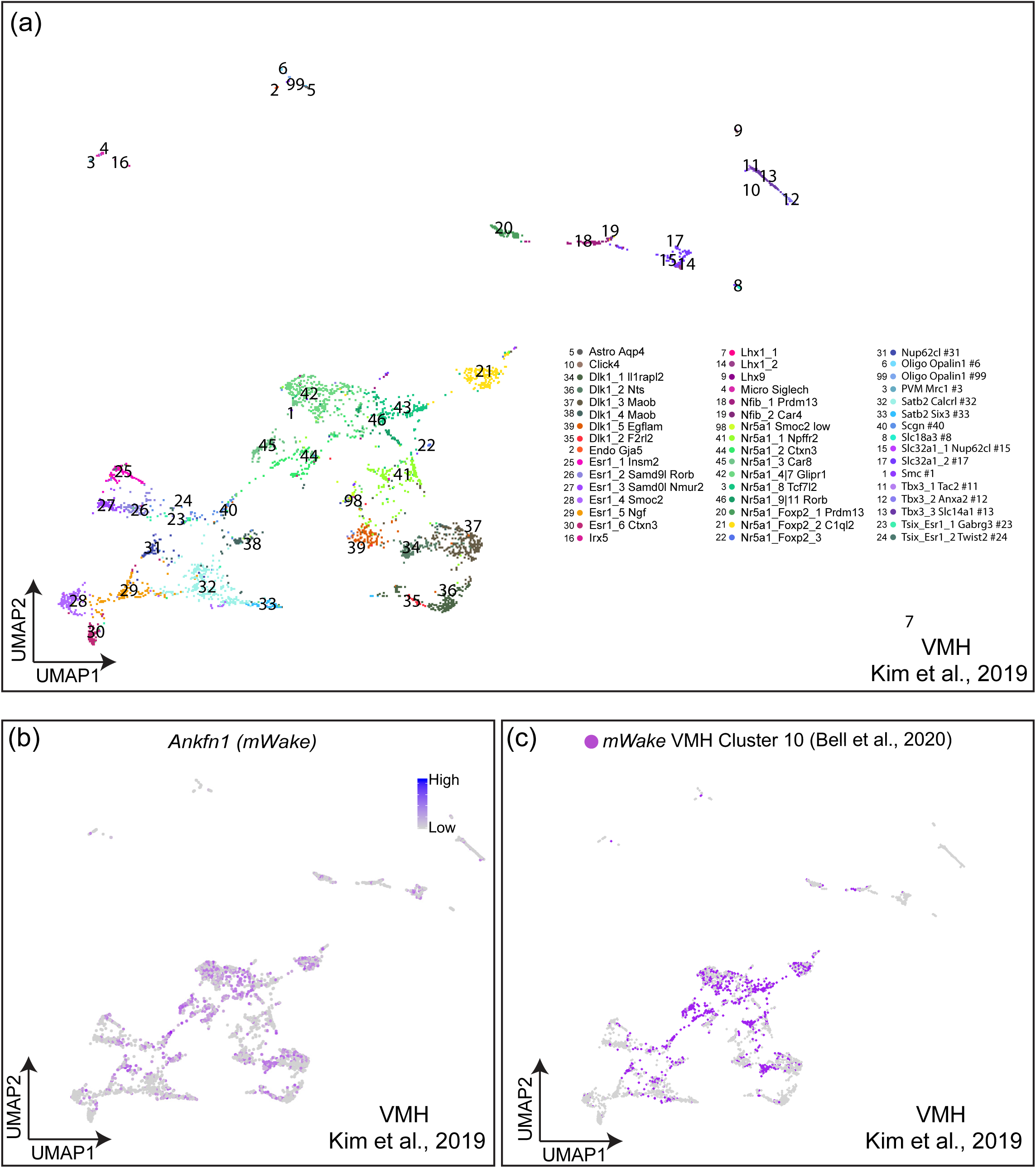
Validation of *mWake* expression identification using key molecular markers. (a) UMAP plot of single-cells isolated from the VMH by Kim *et. al.* (2019). (b) UMAP of same data with expression of *Ankfn1/mWake* in clustered cells shown as color intensity (purple). (c) UMAP plot of same cells with putative *mWake^+^* cells identified by analysis of key molecular markers from *mWake^+^* cells in the VMH of Bell *et. al.* (2020) colored (purple).

**Figure S3.**
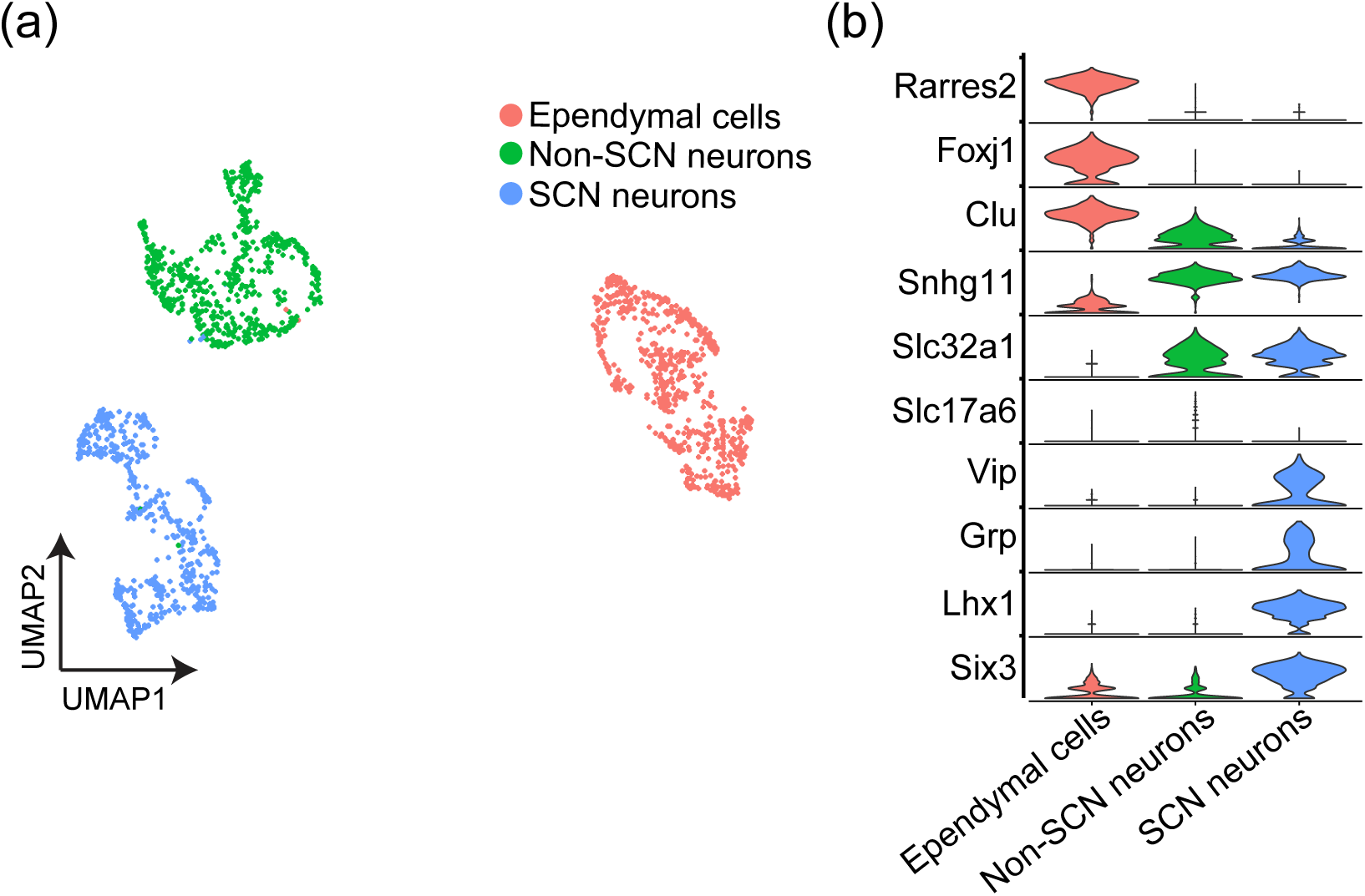
Main scRNA-seq clusters of *mWake^+^* cells from the hypothalamus. (a) UMAP plot of *mWake^+^* scRNA-Seq data from Bell *et. al.* (2020), identifying three main clusters of cells in the hypothalamus: SCN neurons (blue), Non-SCN neurons (green), and ependymal cells (red). (b) Violin plot depicting the expression of key molecular markers within each cluster population. Each row depicts the expression of a single gene, with expression level on the y-axis, and the percentage of the cells expressing at that level forming the width.

**Figure S4.**
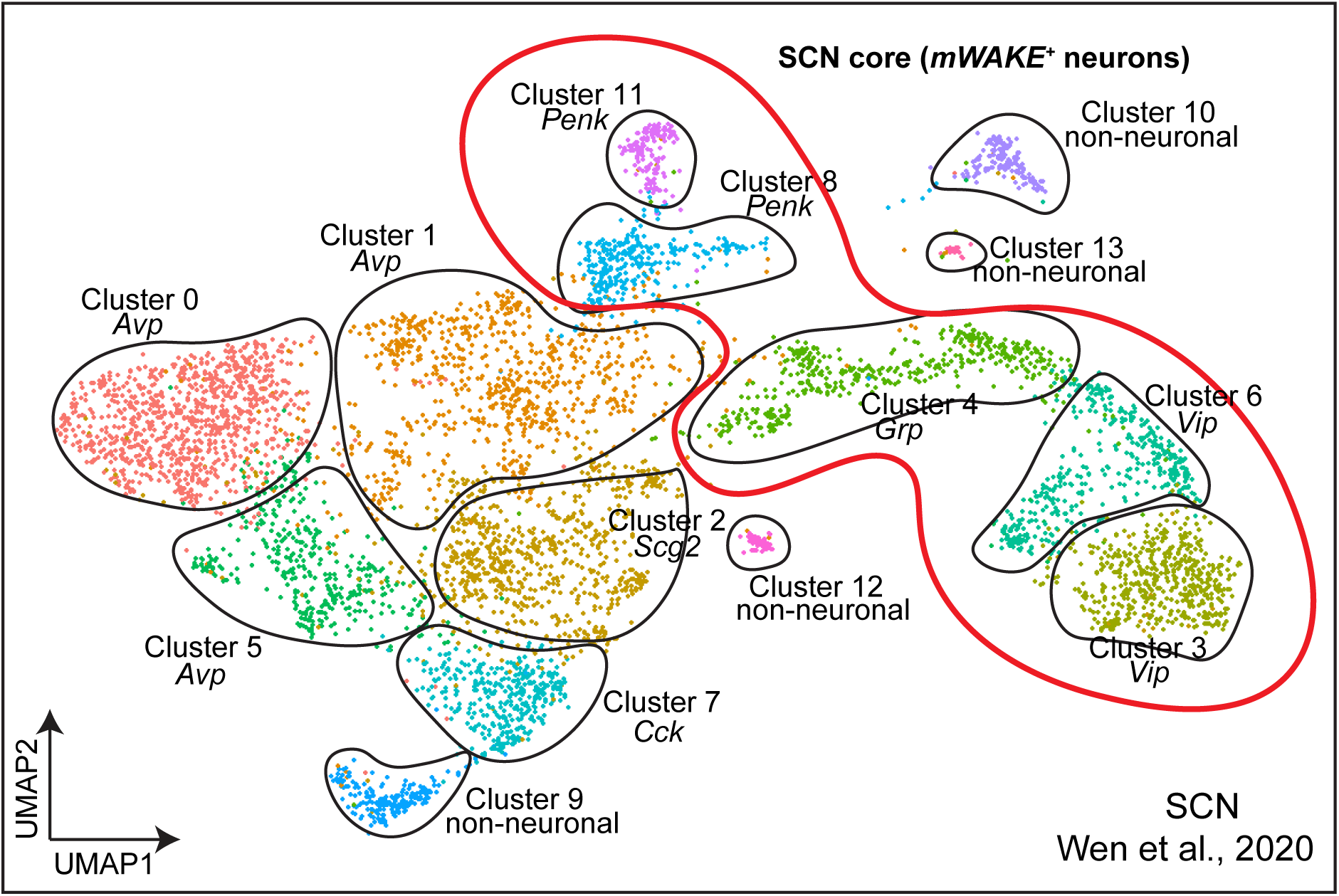
Clustering of SCN scRNA-Seq dataset. (a) UMAP plot of SCN neurons from Wen *et. al.* (2020), with clustering redrawn and labeled with key molecular identities. The thick red line outlines the clusters which are defined by markers of the SCNc.

